# Improving Cardiac Resilience to Ischemia/Reperfusion: The Role of Butyrate in Mitochondrial and Metabolic Recovery

**DOI:** 10.1101/2025.06.23.661163

**Authors:** Bhuban Ruidas, William A. Michaud, Sar R. Lindner, Asishana A. Osho, Shannon N. Tessier, Seyed Alireza Rabi

## Abstract

**Background:** Cardiac transplantation is limited by a persistent shortage of donor organs. While hearts donated after circulatory death (DCD) could expand donors pool, their use is hindered by high rates of primary graft dysfunction (PGD) due to ischemia/reperfusion (I/R)-induced metabolic injury. Here, we investigate the therapeutic potential of the short-chain fatty acid butyrate (BT) to restore metabolic function in cardiomyocytes following I/R.

**Method:** Adult human ventricular cardiomyocytes were used for *in vitro* cell perfusion (IVCP). Following a period of warm ischemia, butyrate (BT) was introduced into a prechilled UW solution to mimic the standard *in situ* cold flush performed during heart explant. Cells were then reperfused with cultural media for 1h at 37°C, after which both cells and perfusate were harvested for spectrometric metabolite profiling and molecular analyses using standard methods.

**Results:** BT reprograms cardiac substrate use from glucose to BT, enhancing mitochondrial oxidative phosphorylation and ATP production, as evidenced by increased lactate clearance and upregulated mitochondrial BT-oxidation enzymes. This metabolic shift restores redox balance by elevating NAD+/NADPH pool and reducing ADP/ATP ratio, while suppressing histone deacetylation and promoting gene expression linked to mitochondrial biogenesis, damage repair, recycling, and turnover *via* enhanced mitofusion and PINK1/Parkin-mediated mitophagy. BT subsequently reduces mitochondrial ROS, enhances electron transport chain activity, and preserves oxidative phosphorylation, thereby lowering caspase-3/7 activity, preventing apoptosis, and promoting cardiomyocyte metabolic recovery.

**Conclusion:** BT restores mitochondrial and metabolic function, preserves ATP synthesis after I/R injury, and mitigates metabolic maladaptation, offering strong potential to improve cardiac viability, graft function, and transplantation outcomes.

**Graphical abstract:** Improving Cardiac Resilience to Ischemia/Reperfusion: The Role of Butyrate in Mitochondrial and Metabolic Recovery

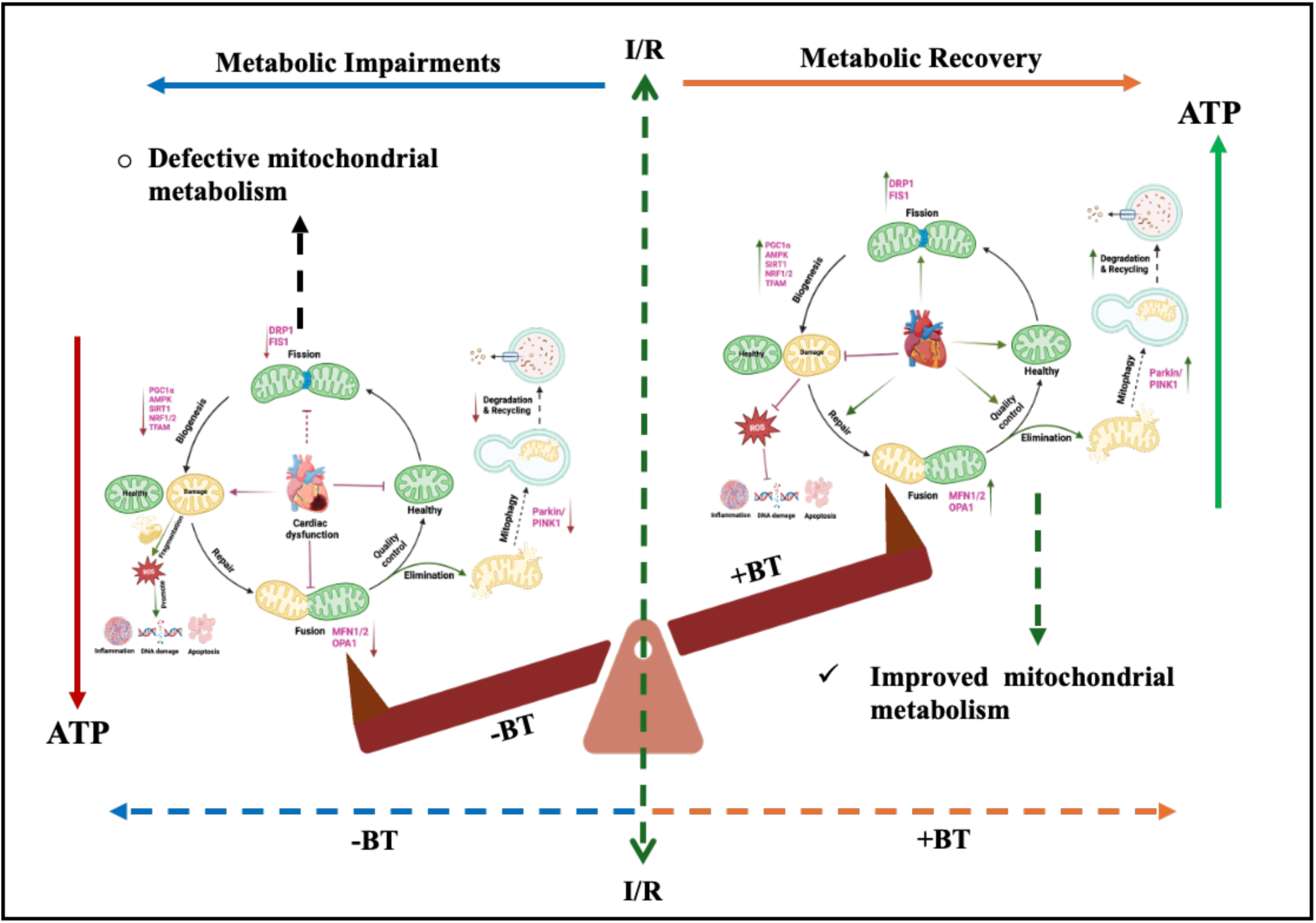

## 1. Introduction

For patients with advanced heart failure and severe symptoms, heart transplantation remains the gold-standard treatment, offering significant improvements in quality of life, functional status, and survival^1^. However, the availability of donor hearts remains critically limited, with only 25-30% of waitlisted patients receiving transplants each year^2^. This shortage, combined with the risk of primary graft dysfunction (PGD), a leading cause of early graft failure and mortality, poses significant challenges^3^. PGD affects approximately 15-41% of transplanted hearts and is frequently attributed to ischemia-reperfusion injury (IRI) incurred during organ procurement and reperfusion^4^. Taken together, the identification of new therapeutic strategies to mitigate IRI is essential for increasing donor organ utilization and addressing the ongoing organ shortage.

Throughout the transplantation process, organs are exposed to IRI at multiple stages. For example, traditional static cold storage (SCS), which is standard for donor’s hearts, can exacerbate ischemic injury, thereby limiting its utility for DCD hearts. Recently, the emergence of *ex vivo* heart perfusion (EVHP), has offered new opportunities to overcome the risk of IRI, especially for DCD hearts^5,6^. In contrast to SCS, warm perfusion technologies provide a continuous oxygen and nutrient supply, while enabling delivery of cardioprotective agents and viability assessment before implantation^7,8^. In this context, optimally managed donor hearts have been shown to recover cardiac function within 24-48 hours^9,10^, although prolonged periods of EVHP are associated with poorer outcomes in DCD heart recipients. Our recent findings demonstrate that with the currently used clinical EVHP system, the Organ Care System (OCS, TransMedics Inc.), time-dependent metabolic changes contribute to progressive cardiac dysfunction, thereby highlighting the importance of addressing overall metabolic health during EVHP is crucial to improvise transplant function and viability^11–13^.

Careful consideration of the metabolic status is especially critical for heart, a highly metabolic organ that demands a substantial amount of energy to sustain spontaneous rhythmic function^14^. The heart’s exceptional metabolic demand is predominantly sustained by mitochondrial oxidative phosphorylation, which is nearly 20 times more efficient than glycolysis (producing 38 ATP vs. 2 ATP per glucose molecule)^15^. As a metabolic omnivore, the adult heart derives ∼95% of its ATP from oxidative pathways: roughly ∼65% from fatty acid oxidation (long-, medium and short chain), ∼ 30% from pyruvate oxidation (via or glycolysis or lactate), and the remaining ∼5% from anaerobic metabolism^16^. Notably, mitochondria, occupying about 25% of the heart’s volume, the highest density among all organs serve as central energy hubs to support its relentless function^17^. Any defects in these mitochondrial oxidative pathways lead to contractile dysfunction and heart failure (HF)^18,19^. Studies also suggest that mitochondrial dysfunction in HF and cardiovascular disease (CVD) involves not only diminished ATP production but also broader maladaptation in mitochondrial structure and function^20–22^. This dysfunction opens new possibilities for mitochondria-targeted therapies to address the diverse roles of mitochondria play in cellular health and CVD^23–25^. Additionally, HF with preserved ejection fraction (HFpEF) is now as prevalent as HF with reduced ejection fraction (HFrEF) in the U.S., both having similar prognoses. HFpEF is increasingly linked to metabolic abnormalities, including obesity and diabetes, suggesting distinct metabolic changes in the heart^26^.

Moreover, FAs depletion has been shown to exacerbate myocardial dysfunction in failing hearts, underscoring the importance of both glucose and FA metabolism^27,28^. In advanced HF, glucose oxidation is downregulated, suggesting a metabolic shift toward alternative substrates, particularly FAs. The reliance on long-chain and medium-chain FAs, which require specific transporters, has been linked to cardiac dysfunction when these transporters are downregulated, a well-established hypothesis in myocardial dysfunction^29^. Evidence also suggests that energy deficiency in the failing heart is due to diminished mitochondrial capacity to oxidize long-chain fatty acids and carbohydrates, leading to reduced ATP production^30,31^. Notably, we have previously shown that human and porcine DCD hearts subjected to EVHP experience rapid depletion of short chain fatty acids due to its inability in utilization of transporter-dependent long-and medium chain fatty acids^32^.

Of note, short-chain fatty acids (SCFAs), with carbon chains of up to six atoms, are crucial for energy metabolism, supplying about 10% of the energy needs in healthy adults^33^. Beyond their role as energy sources, SCFAs function as metabolic regulators with therapeutic potential, as their reduction disrupts energy homeostasis in heart failure (HF) and cardiovascular disease (CVD), making them promising targets for intervention^34,35^. Among SCFAs, butyrate stands out for its superior mitochondrial oxidation efficiency compared to acetate and propionate, making it a highly effective energy substrate^36^. In *ex vivo* Langendorff heart models, supplementation with ketones and butyrate improved myocardial function in heart failure, with butyrate generating more ATP than ketone bodies, further emphasizing its promise as a superior metabolic therapy^36^. Besides, Hypertensive patients also exhibit reduced levels of butyrate-producing bacteria in their gut microbiomes, resulting in decreased serum butyrate levels, underscoring the need to restore this critical metabolic regulator^37^. Given these challenges, short-chain FAs, specifically BT has gained significant clinical interest in metabolic therapies due to their transporter-independent nature and their potential to bypass the limitations of long- and medium-chain FA uptake, offering a more readily available energy source ^38–40^. To date, no studies have assessed butyrate as a metabolic therapy for under-utilized donor hearts. Evaluating its impact on underperforming DCD grafts could reveal ways to preserve metabolism during ischemia/reperfusion, reduce primary graft dysfunction, and expand the pool of transplantable organs.

Herein, we introduce a novel *in vitro* cell-based perfusion (IVCP) platform using human left-ventricular cardiac myocytes that recapitulates the clinical DCD heart procurement sequence. In this model, primary cardiomyocytes undergo successive phase of warm ischemia, followed by cold ischemia to mimic the *in-situ* flush and instrumentation. Finally, cells are cultured at 37°C to mirror normothermic machine perfusion. The primary objective of this study is to administer BT to cardiomyocytes during the cold flush phase, when metabolic and mitochondrial injury is at its peak due to oxygen and nutrients deprivation. Upon reperfusion, this initial insult is compounded by a surge of reactive oxygen species (ROS) and inflammatory mediators that triggers tissue damage and apoptosis. We then evaluate whether BT can preserve mitochondrial bioenergetics and overall metabolic function during the subsequent normothermic perfusion. We hypothesize that BT, by acting as a readily utilizable alternative fuel, will effectively preserve mitochondrial integrity, sustain oxidative metabolism and ultimately promote cardiomyocyte viability and functional recovery following ischemic injury.

## 2. Materials and methods

### 2.1. Primary cell lines and experimental methods

Human cardiac myocytes, HCM and AC10 isolated from adult ventricles were procured from PromoCell (**#12810**) and ATCC (**#CRL-3569**) respectively. HCM were grown in myocyte growth medium (PromoCell C-22070) and AC10 cells were grown in Dulbecco’s Modified Eagle Medium/Nutrient Mixture F-12, DMEM-F12 (ATCC 30-2006) with 12.5% FBS (ATCC 30-2020), 1X Insulin-Transferrin-Selenium (ITS -G) and 2% Horse Serum (Gibco Catalog # 16050130).

2.2 *In vitro cardiac myocytes perfusion and molecular analysis*

Our study aligns with the established human DCD heart transplantation methodology and the detailed heart procurement and perfusion process has been discussed in **supporting information**. Our observations indicate that most underutilized hearts exhibit severe depletion of energy and metabolic substrates, including fatty acids, amino acids, and ketones, highlighting the need for immediate therapeutic intervention^32^. To address this, we introduced a metabolic therapy in an *in vitro* perfusion (IVCP) model of cardiomyocyte (primary cardiac myocytes extracted from the left ventricles of adult human hearts), aligning with the EVHP methods used for DCD hearts. To replicate the process of DCD heart procurement and normothermic machine perfusion, we developed an *in vitro*, primary cardiac cell-based model as shown in **Figure 1A**. BT is introduced during the cold ischemic phase in the cell culture at 4-6°C with 1% O2 for 45-60min of incubation. Briefly, cardiac myocytes cells are grown to be confluent and kept in hypoxic chamber (1% O2) with 5% CO2 at 37°C for 60min of incubation (warm ischemia time; WIT). Following this, immediately cold ischemia to cardiomyocytes were introduced replacing cell media with the pre-chilled University Wisconsin (UW) solutions at 4°C, specified for organ storage and executed targeted metabolic therapy keeping positive, negative and untreated control for 60min of incubation in hypoxic chamber (1% O2) with 5% CO2 at 4-6°C (cold ischemia time; CIT). Next, we performed normoxic perfusion with normal cell media replacing UW solutions (warm at room temperature) at 37°C with 20% O2 and 5% CO2 for 60min of incubation. At each step of this experiment, we collected cell perfusate for electrometabolites studies and cells for detailed molecular and metabolomics analysis.

**Figure 1.**
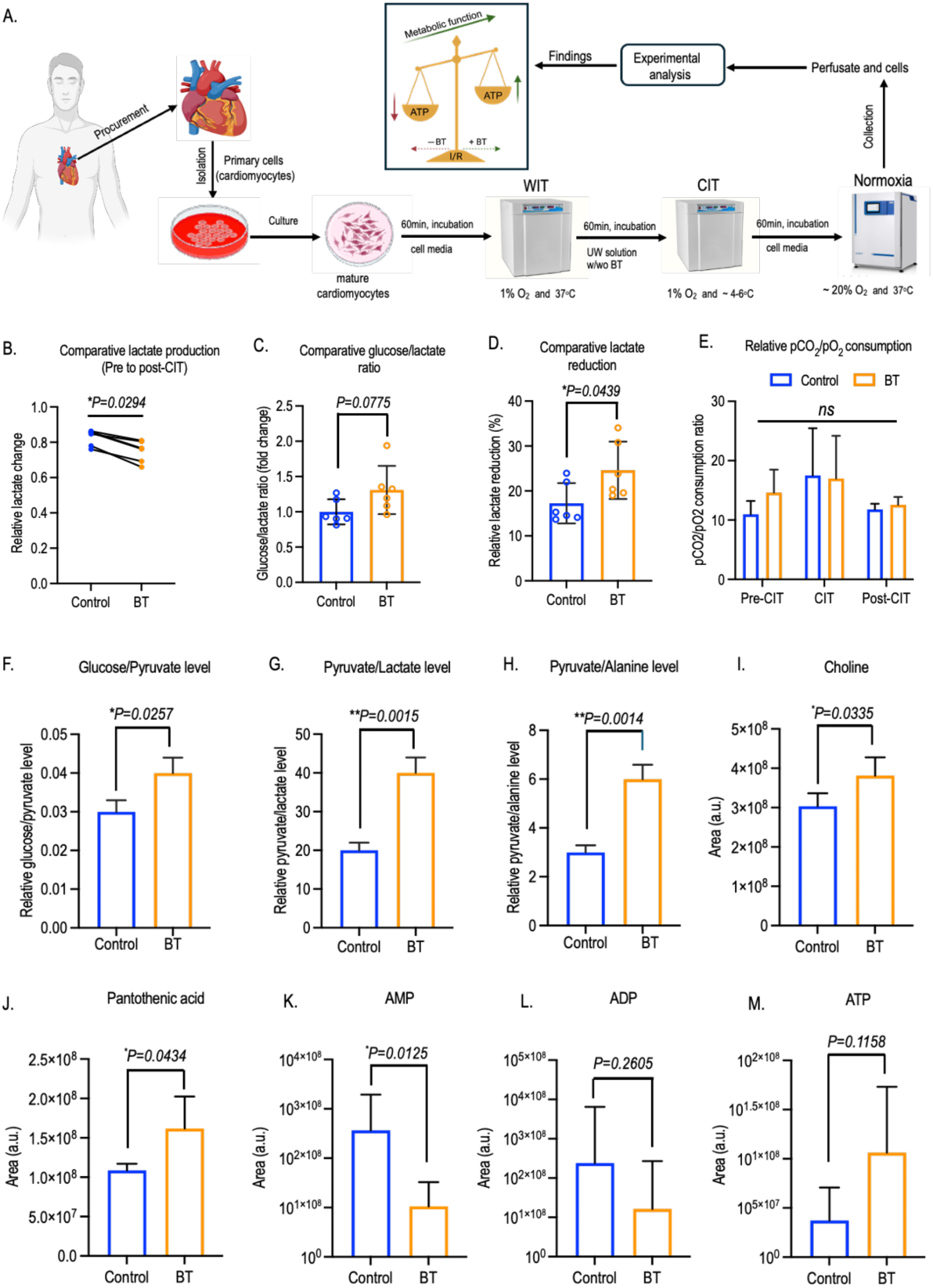
BT upregulates metabolic function in ischemic cardiac myocyte. (**A**) Schematic method of primary cardiac myocytes-based IVCP with BT treatment during CIT in UW solutions followed by normothermic perfusion with cell-based media improves metabolic function compared to untreated control (UC). In each step of perfusion, cell perfusate were collected whereas cells were collected at the end of normothermic perfusion before spectrometric metabolomic and molecular analysis. Changes in cell perfusate-based electrometabolites includes-comparative lactate production (**B**), glucose/lactate ratio (**C)**, lactate reduction (**D**) and relative consumption of pCO_2_/pO_2_ (**E**) are depicted. Cell-based metabolomes profiles with significant changes in the level of Glucose/Pyruvate (**F)**, Pyruvate/Lactate (**G**), Pyruvate/Alanine (**H**) ratio are shown quantitively. Cell-based other nutrients include-choline (**I**) and Pantothenic acid or Vitamin B5 has been depicted and quantified. Changes in high energy phosphate includes AMP (**J**), ADP (**K**) and ATP (**L**) has been shown and analyzed. Data from all BT treatments are quantified and plotted based on results from three to five independent experiments, and comparisons are made against the control group. The bar represents the mean ± SD (standard deviation). Statistical analyses are conducted using GraphPad Prism Version 10.3.1 (464), and significance level are denoted as \**p*<0.05, \*\**p*<0.01, based on Student’s t-test. IVCP; *In vitro* cell perfusion, WIT; Warm ischemia time, CIT; Cold ischemia time, pCO_2_; Partial carbon dioxide, pO_2_; Partial O_2_, AMP; Adenosine monophosphate, ADP; Adenosine diphosphate, ATP; Adenosine triphosphate.

### 2.3 Perfusate-based electrometabolites and cell-based metabolomics assay

To analyze perfusate-derived electrometabolites and cell-based metabolomics, 5×10^4^ cells were seeded in 12-well plate and grown to confluence before performing IVCP, with or without BT treatment. For electrometabolite analysis, perfusate samples were collected at different stages of IVCP: from baseline hypoxia, during cold ischemia with UW solution, and during warm ischemia with cell media and stored at -80°C before recording the electrometabolites profiles using the Siemen’s RAPIDPoint ®500 systems. All the data were quantified relative to untreated control. For cell-based metabolomics analysis, cells were trypsinzed and pelleted down to discard the cell lysate. Next the cell pellets were washed with 1X PBS and re-pelleted to store in mass spectrometry grade pre-chilled methanol at -80°C prior to dropping it at Harvard center of mass spectrometry core facility. Detailed materials and method of cell and molecular analyses are described in the **supplementary information**.

### 2.4 Statistical analysis

All experiments were performed with a n = 3-7 independent replicates. Data are presented as the mean ± standard error of mean (s.e.m) and standard deviation (SD). MS results were correlated standard calibration samples and only results with high confidence level of >90% were included in the analysis. Statistical significance was analyzed using as unpaired two-tailed Student’s t-test in Prism (GraphPad Software, La Jolla California USA). Mitochondrial distributions are quantified by Pearson’s linear correlation (Two-tailed) analyses the 95% of confidence interval in figure 4D. Statistical significance thresholds were set at p<0.05 (*), p<0.01 (**), p<0.001(***), and p<0.0001(****).

## 3. Results

### 3.1. BT upregulates metabolic function in ischemic cardiac myocyte

Using a cell viability assay combined with ADP/ATP ratio quantification at varying BT concentrations, we determined that a 1 mM dose of BT maintained over 90% cell viability while promoting ahigh rate of ATP generation (Figure S1A-S1C). Consequently, this concentration was used for all subsequent experiments described in the manuscript.

BT treatment significantly reduced lactate production in the BT-treated cells perfusate before and after CIT followed by elevation in glucose/lactate ratio and lactate reduction (**Figure 1B-1D**) compared to untreated control. However, there was no significant changes in relative pCO2/pO2 consumption rate, electrolytes levels including Na^+^, K^+^, Ca^++^, Cl^-^ and pH (**Figure 1E** & **Figure S1D, S1E**). In cell-based metabolomics studies, BT significantly improves glucose/pyruvate ratio and reduces lactate and alanine production in response to pyruvate consumption (**Figure 1F-H**). Additionally, BT significantly improve free glucose level but there were no significant changes in pyruvate level, whereas a significant decline in lactate and trends to be decline in alanine levels were found compared to untreated control (**Figure S1F-S1J & Table S1**). Moreover, BT effectively elevates levels of α-ketoglutarate, a key metabolic intermediate of the tricarboxylic acid (TCA) cycle, which is shown to have a cardioprotective effects (**Figure S1K**)^41^. BT-treated cardiac cells also exhibit increased levels of other nutrients, including choline and pantothenic acid, alongside a reduction in phosphocholine (**Figure 1I, 1J & Figure S1L**). Choline, beyond its role in lipid transport and metabolism, has been shown to alleviate cardiac dysfunction by modulating key proteins involved in ketone body and fatty acid metabolism^42^. Pantothenic acid (vitamin B5), essential for synthesizing coenzyme A, plays a crucial role in fatty acid synthesis and degradation. Importantly, phosphocholine is produced from choline in an ATP-dependent reaction catalyzed by choline kinase, produces ADP as byproduct. Under ischemic stress, cardiac cells effectively utilize BT to restore metabolic function, leading to decreased phosphocholine and ADP levels and increased choline and ATP production. This restoration is confirmed by reduced AMP and ADP levels alongside elevated ATP levels (**Figure 1K-1M**). These findings are reinforced by a detailed quantification of BT’s molecular utilization and ATP synthesis, confirming its role in supporting cardiac cell’s metabolism under ischemic stress.

### 3.2. BT fuels cardiomyocytes by enhancing mitochondrial oxidation and ATP production

Metabolome profiling of BT-treated cardiac cells shows a marked increase in glucose levels without a corresponding rise in pyruvate, suggesting a reduction in glucose oxidation and a metabolic shift toward BT as an alternative energy source. BT readily diffuses into the mitochondrial matrix before broken down enzymatically to fulfill energy demands in ischemic cardiac cells, where it is first converted to butyryl-CoA by acyl-CoA synthetase medium-chain family member 3 (ACSM3). It then undergoes sequential breakdown into crotonyl-CoA, 3-hydroxybutyryl-CoA, acetoacetyl-CoA, and finally acetyl-CoA, catalyzed by short-chain acyl-CoA dehydrogenase 1 (SCAD1), Short-chain enoyl-CoA hydratase 1 (ECHS1), 3-Hydroxyacyl-CoA dehydrogenase (HADH), and acetyl-CoA acetyltransferase 1 (ACAT1), respectively (**Figure 2A**). A marked increase in BT metabolism-associated catalytic enzymes include-ACSM3, SCAD1, ECHS1, HADH and ACAT1 confirms the successive utilization of BT for metabolic function and ATP synthesis (**Figure 2B-2E**, **S2A & Table S2**). However, expressions of long chain fatty acid (LCFA) transporter, FAT/CD36 and medium chain fatty acid transporter, MCT1 were unchanged (**Figure 2F** & **2G**). Additionally, BT enhances the expression of key mitochondrial proteins involved in fatty acid metabolism, such as carnitine palmitoyltransferase 1 and 2 (CPT1 and CPT2), with CPT1 showing a trend towards a significant upregulation (**Figure 2H** & **S2B**). Furthermore, BT significantly upregulates the expression of the mitochondrial enzyme pyruvate dehydrogenase kinase 4 (PDK4), a key regulator of cellular energy homeostasis and glucose metabolism (**Figure 2I**). The increased expression of PDK4 supports the enhanced reliance of cardiac myocytes on BT metabolism during I/R, emphasizing a successive metabolic shift from glucose to BT utilization^43^. In contrast, the levels of pyruvate dehydrogenase (PDH), lactate dehydrogenase A (LDHA), and alanine aminotransferase (ALAT)-enzymes involved in the conversion of pyruvate to acetyl-CoA, lactate, and alanine respectively remained unchanged (**Figure S2C-2E**). Moreover, BT treatment acts as a metabolic regulator by enhancing NAD^+^/NADH ratio in stressed cardiac cells, which stimulate ATP synthesis machinery^44^. This increase in ATP synthesis is evidenced by both qualitative and quantitative elevations in ATP-probe intensities in cardiac cells under stress, compared to untreated control cells (**Figure 2J-2L**). This enhancement in ATP production is further elucidated by the subsequent quantification of mitochondrial respiratory complexes, which provides insights into the underlying mechanisms driving this metabolic shift for readily available energy sources.

**Figure 2.**
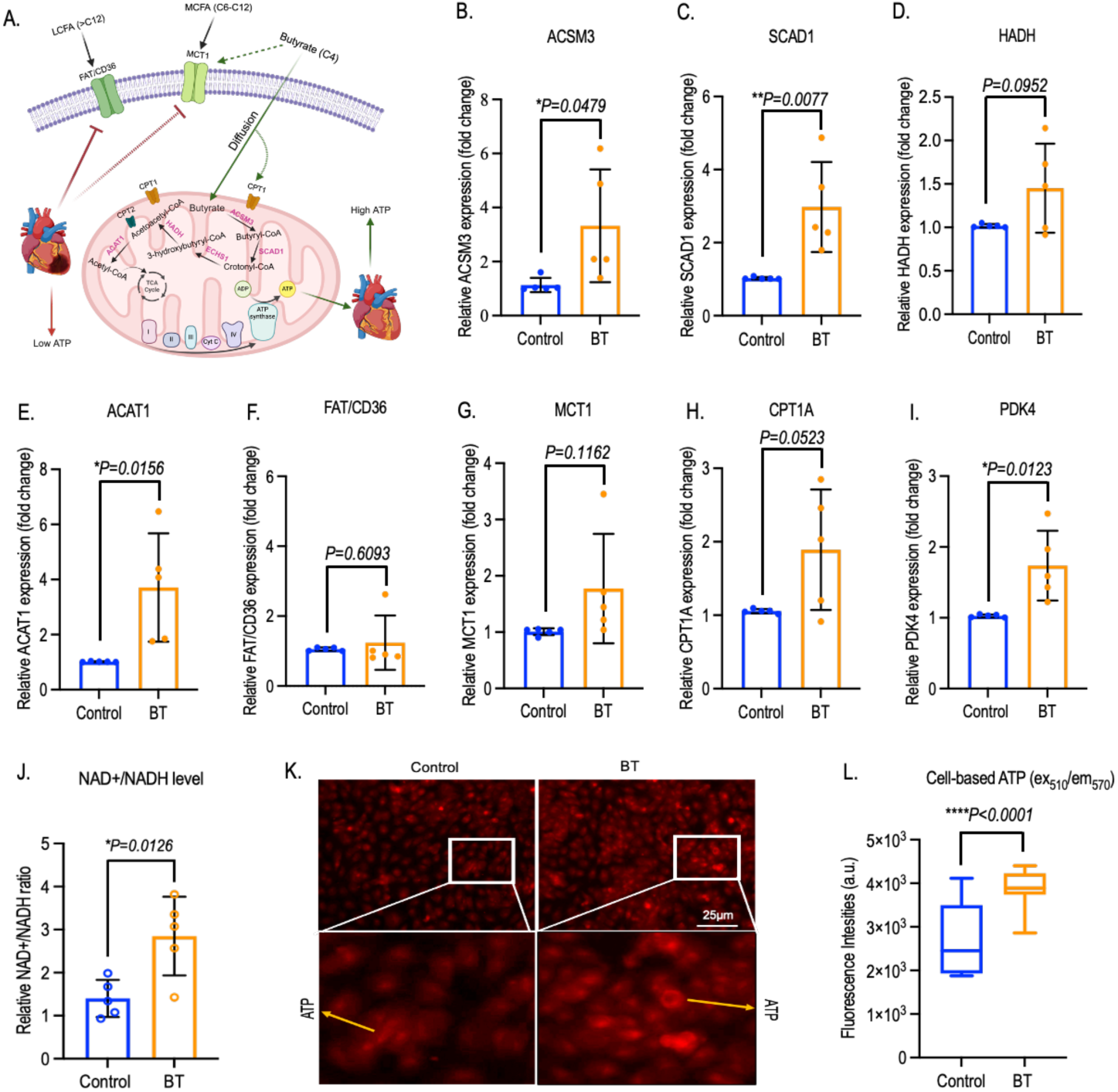
BT fuels cardiac cells by enhancing mitochondrial oxidation and ATP production. **(A)** A detailed method of BT utilization in the cardiac cell’s inner mitochondrial matrix as readily available energy resource are depicted schematically in response to cardiac function and dysfunction and all genes associated with BT metabolism were quantified by qRT-PCR analysis. Relative improvements in the genes associated with sequential breakdown of BT into Acetyl-CoA-includes ACSM3 **(B)** converts BT to butyryl-CoA, SCAD1 **(C)** converts butyryl-CoA to crotonyl-CoA, HADH (**D**) converts 3-hydroxybutyryl-CoA to acetoacetyl-CoA and finally ACAT1 **(E)** converts acetoacetyl-CoA to acetyl-CoA has been depicted. Expression of long chain fatty acid transporter gene, FAT/CD36 **(F)** and medium chain transporter gene, MCT1 **(G)** expression has been analyzed and depicted. Expression of outer mitochondrial fat-transporter protein, CPT1A **(H)** and, pyruvate to acetyl-CoA conversion inhibitor gene PDK4 **(I)** and are depicted. Comparative NAD^+^/NADH levels **(J)** are shown followed by the depiction of intracellular ATP imaging **(K)** using fluorescence probe and real time measurements of cell-based fluorescence at excitation and emission wavelength of 510nm and 570nm respectively. Data from all BT treatments are quantified and plotted based on results from three to five independent experiments, and comparisons are made against the control group. Each dot in gene expression and NAD^+^/NADH ratio graph represents an independent experiment, with the mean value derived from three identical replicates. Bar represents the mean ± SD (standard deviation). Statistical analyses are conducted using GraphPad Prism Version 10.3.1 (464), and significance level are denoted as \**p*<0.05, \*\**p*<0.01, \*\*\*\**p*<0.0001, based on Student’s t-test. qRT-PCR; Quantitative reverse transcription polymerase chain reaction, ACSM3; Acyl-CoA synthetase medium-chain family member 3, SCAD1; Short-chain acyl-CoA dehydrogenase 1, ECHS1; Short-chain enoyl-CoA hydratase 1, HADH; 3-Hydroxyacyl-CoA dehydrogenase; ACAT1; Acetyl-CoA acetyltransferase 1; FAT/CD36; fatty acid translocase/cluster of differentiation 36, MCT1; fatty acid medium chain transporter, PDK4; pyruvate dehydrogenase kinase 4, CPT1A; Carnitine palmitoyltransferase I, CPT2; Carnitine palmitoyltransferase 2, NAD; nicotinamide adenine dinucleotide, NADH; nicotinamide adenine dinucleotide hydrogen.

### 3.3. BT improves ETC complex activities and restore ATP supply under I/R-induced stress

The mitochondrial electron transport chain (ETC) plays a vital role in generating the energy required for proper heart function, and any defects within this complex can lead to metabolic dysfunction (**Figure 3A**). A significant increase in the gene expressions-associated with mitochondrial complex activities confirms the functional activities in cardiac cells under ischemic stress (**Figure 3B** & **Table S3**). Furthermore, BT upregulates the expression of genes associated with mitochondrial Complex I, including NDUFA1, NDUFA4, NDUFB3, NDUFS2, MT-ND1, MT-ND2, MT-ND3, MT-ND4, MT-ND4L, MT-ND5, and MT-ND6. These genes play crucial roles not only in mitochondrial bioenergetics but also in cardiac protection, extending beyond their primary function of electron transfer from NADH to ubiquinone. All complex I-related genes show a significant increase in expression, with NDUFA1 exhibiting a trend toward significance (**Figure 3C-3K**, & **Figure S3A, S3B**). BT also upregulates the expression of genes associated with Complex II, including SDHA, SDHB, and SDHD, which primarily facilitate electron transfer from succinate to ubiquinone within the ETC complex (**Figure 3L, 3M** & **Figure S3C**). Additionally, BT enhances the expression of genes involved in Complex III, which receives electrons from both Complex I and Complex II and maintains the proton gradient that drives ATP synthesis at Complex V. BT upregulates Complex III-associated genes, including CYC1, CYTB, and UQCRQ, with CYC1 and CYTB showing significantly high expression levels (**Figure 3N, 3O** & **Figure S3D**). BT also upregulates the expressions of genes including COX1, COX2, COX3, COX5A and COX7A1-associated with Complex IV, the final electron acceptor and a major regulator of oxidative phosphorylation in the ETC complex. Expressions of COX1, COX3 and COX5A genes found to be highly significant, whereas the expressions of COX2 and COX7A1 genes trends to be significant with BT treatment compared to untreated control (**Figure 3P-3Q** & **Figure S3E, S3F**). Finally, BT effectively enhances the activity of the highly conserved Complex V enzyme, ATP synthase, which is essential for the energy-consuming processes of cardiac contraction and relaxation. ATP synthase catalyzes the synthesis of ATP from ADP at its F1 sector, the enzyme’s catalytic site. BT significantly upregulates genes associated with ATP synthase activity, including MT-ATP6, MT-ATP8, ATP5A1, and ATP5D, while ATP5E shows a trend toward significance compared to the untreated control (**Figure 3S, 3T** & **Figure S3G-S3I**). In real-time measurements of mitochondrial respiratory Complex I and Complex II activity in live cells, BT markedly enhances the activity of both complexes, indicating a potential restoration of ATP synthesis in cardiac cells under ischemic stress (**Figure 3U** & **3V**). This effect is further confirmed by a quantifiable improvement in the cellular ADP/ATP ratio (**Figure 3W**). Comparative profiling of the ADP/ATP ratio in cardiac cells treated with BT, long-chain fatty acid (palmitate), a lipid mixture, and resveratrol (as a negative control) further highlights the superior efficacy of BT as a metabolic therapy (**Figure S3J**). The reduction in the cellular ADP/ATP ratio with BT treatment confirms an increase in ATP synthesis, likely resulting from sustained mitochondrial oxidative respiration and improved mitochondrial function.

**Figure 3.**
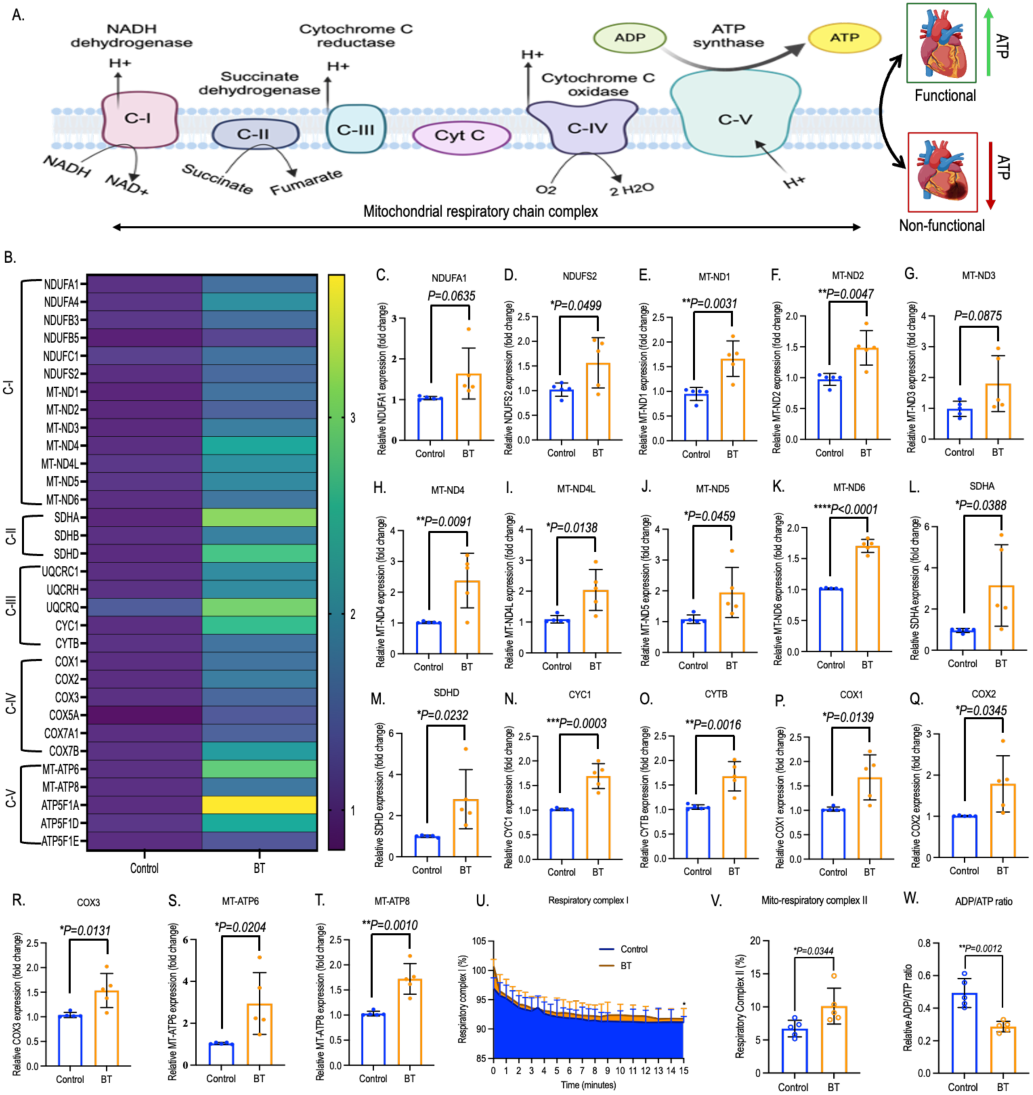
BT improves ETC complex activities and restore ATP supply under I/R-induced stress in cardiac myocytes. **(A)** A schematic representation of mammalian respiratory chain complex induces ATP synthesis and its associative impact on functional and non-functional hearts has been depicted and all genes associated with ETC complex were quantified by qRT-PCR analysis. **(B)** A comparative heat-map of ETC complex-associated genes in perfused cardiac myocytes with or without BT has been depicted. Significant changes in complex-I associated genes includes-NDUFA1 **(C)**, NDUFS2 **(D)**, MT-ND1 **(E)**, MT-ND2 **(F)**, MT-ND3 **(G)**, MT-ND4 **(H)**, MT-ND4L **(I)**, MT-ND5 **(J)** and MT-ND6 **(K)** are depicted. Significant changes in complex-II associated genes-SDHA **(L)** and SDHD **(M)** are shown quantitively. Significant changes in complex-III associated genes-CYC1 **(N)** and CYTB **(O)** are depicted. Significant changes in complex-IV associated genes-COX1 **(P)**, COX2 **(Q)** and COX3 **(R)** are shown quantitively. Significant changes in complex-V associated genes MT-ATP6 **(S)** and MT-ATP8 **(T)** are depicted quantitively. Comparative quantification of overall cell-based mitochondrial respiratory complex I **(U)** and complex II **(V)** activities are depicted quantitively. Significant changes in ADP/ATP ratio **(W)** exhibiting metabolic coupling of mitochondrial OXPHOS are depicted and quantified. Data from all BT treatments are quantified and plotted based on results from five to six independent experiments, and comparisons are made against the control group. Each dot in gene expression, respiratory complex and ADP/ATP ratio graph represents an independent experiment, with the mean value derived from three identical replicates for gene expressions and five identical replicates for rest of the analyses. Bar represents the mean ± SD (standard deviation). Statistical analyses are conducted using GraphPad Prism Version 10.3.1 (464), and significance level are denoted as \**p*<0.05, \*\**p*<0.01, \*\*\**p*<.001, \*\*\*\**p*<0.0001, based on Student’s t-test. qRT-PCR; Quantitative reverse transcription polymerase chain reaction, NDUFA1 and NDUFA4; NADH dehydrogenase (ubiquinone) 1 alpha subcomplex 1 and 4, NDUB3 and NDUB5; NADH dehydrogenase (ubiquinone) 1 beta subcomplex 3 and 5, NDUFC1; NADH dehydrogenase [ubiquinone] 1 subunit C1, NDUFS2; NADH dehydrogenase [ubiquinone] iron-sulfur protein 2. MT-ND1; NADH-ubiquinone oxidoreductase chain 1. SDHA; Succinate dehydrogenase [ubiquinone] flavoprotein subunit A, SDHB; Succinate dehydrogenase [ubiquinone] iron-sulfur subunit B, SDHD; Succinate dehydrogenase [ubiquinone] cytochrome b small subunit D, UQCR(C1,H, and Q); Cytochrome b-c1 complex(subunit 1, 6, and 8), CYC1; Cytochrome C1, CYTB; cytochrome b, COX (1, 2, 3, 5A,7A1and 7B); Cytochrome oxidase (subunit 1, 2,3, 5A, 7A1 and 7B), MT-ATP6 and ATP8; Mitochondrially encoded ATP synthase membrane subunit 6 and 8, ATP5F1(A, D and E); ATP synthase F1 (subunit alpha, delta, and epsilon), OXPHOS; Oxidative phosphorylation.

### 3.4. BT resilience I/R-induced mitochondrial injury and enhance its viability

Mitochondrial damage and loss can severely impair oxidative phosphorylation and reduce ATP production in cardiac cells, particularly under ischemic stress^45^. BT efficiently restore the mitochondrial functional integrity through increments in mitochondria number and viability (**Figure 4A**). BT significantly upregulates mitochondrial viability and number in cardiac cells under stress (**Figure 4B & Figure S4A**). BT also improves the mitochondrial structure by length which is trends to be significant while mitochondrial width remained unchanged (**Figure 4C, S4B**). Additionally, mitochondrial distribution by length is improved in BT-treated cardiac cells under stress, resulting enhanced mitochondria health which were further confirmed through the imaging of viable mitochondria with Mito-tracker probe (**Figure 4D, 4E**). Viable mitochondria intensities have significantly elevated in BT-treated cells compared to untreated control, resulting the increment in mitochondria function (**Figure 4F**). Analysis of gene-associated with mitochondrial DNA synthesis has further confirmed the prolong mitochondrial viability under stress. BT enhances the mitochondrial biogenesis by successive inhibition of histone deacetylase which is confirmed through the depletion in histone deacetylase class I (HDAC1) gene expressions (**Figure 4G**). In turn, BT enhances the expressions of mitochondrial DNA replication-associated genes-polymerase gamma 1 and 2 (POLG1 and POLG2), transcription associated gene-mitochondrial transcription factor A (TFAM) and translation-associated gene-mitochondrial translational factor 2 (MTIF2). BT significantly upregulates POLG1, POLG2 and MTIF2 whereas TFAM is trends to be significant compared to untreated control (**Figure 4H-4J, & S4C**). The enhanced mitochondrial biogenesis therefore confirms BT’s potential to restore ATP production and support metabolic recovery, enabling cardiac cells to meet their high energy demands during I/R-induced stress.

**Figure 4.**
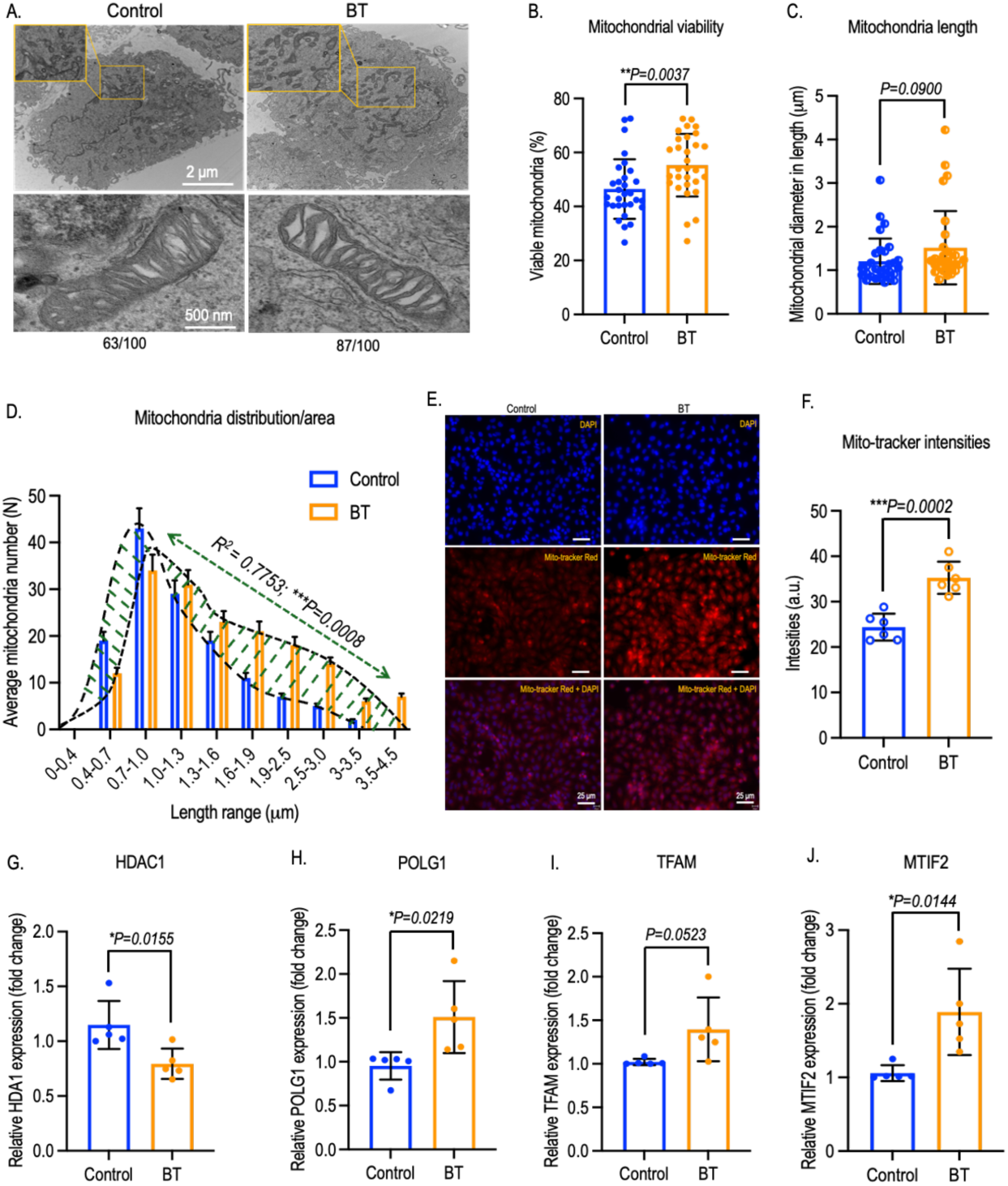
BT resilience I/R-induced mitochondrial injury in cardiac myocytes and improve mitochondrial viability. **(A)** Electron microscopic view of perfused cardiac cell’s mitochondria with or without BT treatment at the end of normothermic perfusion has been depicted using electron microscopy analysis (Scale bar - 2μm and 500nm respectively, Viability number count per area normalized to 100). **(B)** Significant changes in mitochondrial viability percentages and mitochondria length **(C)** are depicted and quantified using ImageJ analysis**. (D)** Distribution of mitochondria by length are depicted and quantified (Dotted green line showing the changes in mitochondria length in perfused cardiac myocytes with or without BT). Mitochondrial distributions were quantified using Pearson’s linear correlation analysis, yielding a 95% confidence interval (0.5631 to 0.9715) and an R² value of 0.7753. This indicates a statistically significant correlation between the control and BT treatment groups**. (E)** Viable mitochondria imaging using Mito-tracker probe followed by quantification of intensities **(F)** are shown quantitively (Scale bar - 25μm). Significant changes in the expression of genes-associated with mitochondrial biogenesis and viability includes-HDAC1 **(G)**, POLG1 **(H)**, TFAM **(I)** and MTIF2 **(J)** has been depicted quantitively using qRT-PCR analysis. Data from all BT treatments are quantified and plotted based on results from five to six independent experiments, and comparisons are made against the control group. Each dot in gene expression graph represents an independent experiment, with the mean value derived from three identical replicates. Bar represents the mean ± SD (standard deviation). Statistical analyses are conducted using GraphPad Prism Version 10.3.1 (464), and significance level are denoted as \**p*<0.05, \*\**p*<0.01, \*\*\*\**p*<0.0001, based on Student’s t-test. qRT-PCR; Quantitative reverse transcription polymerase chain reaction, HDAC1; histone deacetylase class I, POLG1; Polymerase gamma 1, TFAM; Mitochondrial transcription factor A, MTIF2; Mitochondrial translation initiation factor 2.

### 3.5. BT improves mitophagy systems and limits mitochondrial ROS production

Under stress conditions, mitophagy is essential for maintaining mitochondrial quality by selective elimination of damaged mitochondria through the Parkin/PINK1-mediated degradation pathway (**Figure 5A & Table S4**). This protective mechanism helps regulate excessive reactive oxygen species (ROS) production, preventing cellular dysfunction^46^. During ischemic injury in cardiac cells, mitochondrial function in cells is compromised as damaged mitochondria accumulate due to impaired mitophagy systems, leading to elevated ROS production. This increase in ROS disrupts DNA synthesis and triggers apoptosis. The depletion of genes involved in mitochondrial biogenesis has a critical impact on the impairment of the mitophagy system. In this context, BT significantly boosts the expression of key genes associated with mitochondrial bioenergetics, including peroxisome proliferator-activated receptor-γ coactivator 1α (PGC-1α), AMP-activated protein kinase (AMPK), Sirtuin 1 (SIRT1), Nuclear Respiratory Factor 1 (NRF1), and nuclear factor erythroid 2-related factor 2 (NRF2). Beyond their roles in cardiac mitochondrial energy metabolism, PGC-1α and SIRT1 are essential for supporting mitochondrial oxidative phosphorylation and respiration in cardiac muscle. Meanwhile, AMPK serves as a cellular energy sensor, regulating mitochondrial function and overall bioenergetics in the heart. BT effectively upregulates PGC-1α, SIRT1, and AMPK expression, helping to restore mitochondrial bioenergetics in ischemic cardiac cells (**Figure 5B-5D**). Additionally, the transcription factors NRF1 and NRF2 play crucial roles in maintaining mitochondrial homeostasis and function in cardiac cells, enhancing ATP production efficiency. BT significantly upregulates NRF1 and NRF2 activity, helping to restore the energy balance of the mitophagy system and support overall mitochondrial health (**Figure S5A, S5B**). In the mitophagy system, damaged mitochondria are repaired through mitochondrial fusion, a process regulated by proteins such as mitofusin 1 (MFN1), mitofusin 2 (MFN2), and optic atrophy 1 (OPA1). Mitochondrial fission, which enables the production of new mitochondria, is governed by proteins like dynamin-related protein 1 (DRP1) and mitochondrial fission protein 1 (FIS1). Once damaged mitochondria are fused with healthy mitochondria for repair, the remaining defective mitochondria are marked for degradation. Proteins such as Parkin and PINK1 then target these marked mitochondria, facilitating their removal from the mitochondrial matrix. This process helps lower cellular ROS levels and supports overall cellular health. In this context, BT significantly enhances the expression of DRP1 and FIS1, promoting mitochondrial fission and supporting the synthesis of healthy mitochondria (**Figure 5E** & **Figure S5C**). This is followed by an increase in MFN1, MFN2, and OPA1, leading to the successive fusion and restoration of mitochondrial quality (**Figure 5F, 5G** & **Figure S5D**). Notably, MFN2 activity plays a pivotal role in initiating mitophagy through the PINK1/Parkin pathway, as demonstrated by elevated levels of MFN2 and PARK2/Parkin observed in antibody-based cytochemistry analyses (**Figure 5H**). This activation is further supported by increased levels of PINK1 and Parkin, which promote the degradation of damaged mitochondria and reduces excessive ROS production (**Figure 5I** & **5J**). Real-time ROS measurements in BT-treated cardiac cells confirm the significant reduction in mitochondrial ROS, suggesting that BT effectively restores mitophagy systems and enhances mitochondrial quality control (**Figure 5K**). However, no significant changes in hypoxia levels were observed after transitioning from hypoxia to normoxia (**Figure S5E & S5F**). Additionally, the sustained reduction in mitochondrial ROS offers promising potential for regulating apoptotic cell death under ischemic stress, a key factor linked to systolic dysfunction in the heart.

**Figure 5.**
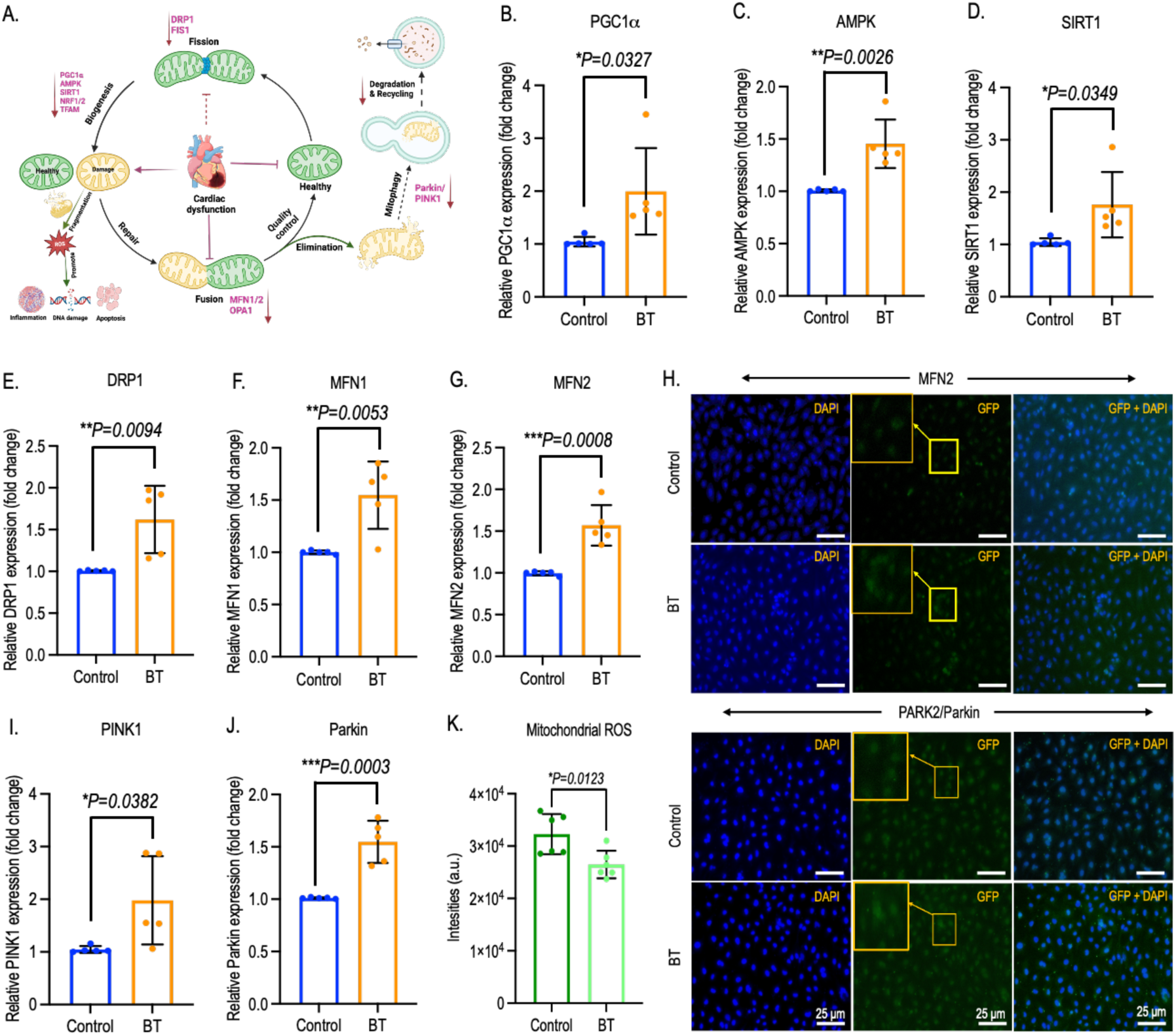
BT improves cardiac cell’s mitophagy systems and limits mitochondrial ROS production. **(A)** A schematic illustration depicting consequences of impaired mitophagy on cardiac health has been depicted and all genes associated with mitophagy system were quantified by qRT-PCR and cytochemistry analysis. Significant elevation in mitochondrial biogenesis associated genes includes-PGC-1α **(B)**, AMPK **(C)** and SIRT1 **(D)** are depicted quantitively. Significant changes in mitochondria dynamics (fission and fusion)-associated genes includes-DRP1**(E)**, MFN1 **(F)** and MFN2 **(G)** are shown respectively. **(H)** Immunocytochemistry with MFN2 and mitophagy protein marker, PARK2/parkin are depicted (Scale bar - 25μm). Mitophagy associated genes includes-Parkin **(I)** and PINK1 **(J)** are depicted comparatively. Significant changes in mitochondrial ROS **(K)** generations in BT-treated perfused cardiac myocytes are depicted quantitively. Data from all BT treatments are quantified and plotted based on results from to five to six independent experiments, and comparisons are made against the control group. Each dot in gene expression and ROS analysis graph represents an independent experiment, with the mean value derived from three identical replicates for gene expressions and five identical replicates for ROS analysis. Bar represents the mean ± SD (standard deviation). Statistical analyses are conducted using GraphPad Prism Version 10.3.1 (464), and significance level are denoted as \**p*<0.05, \*\**p*<0.01, \*\*\**p*<.001, based on Student’s t-test. PGC-1α; Peroxisome proliferator-activated receptor-γ coactivator 1α, AMPK; AMP-activated protein kinase, SIRT1; Sirtuin 1, DRP1; Dynamin-related protein 1, MFN1; Mitofusin 1, MFN2; Mitofusin 2, PINK1; PTEN-induced kinase 1, qRT-PCR; Quantitative reverse transcription polymerase chain reaction, ROS; Reactive oxygen species.

### 3.6. BT effectively limit apoptotic cell death in cardiac myocyte

Both acute and chronic loss of cardiac myocytes due to apoptosis during reperfusion injury is well established^47^. Caspase 8 and caspase 9 play key roles in initiating early apoptosis, while caspase 3, caspase 6, and caspase 7 drive the progression to intermediate and late stages of apoptosis in cardiac cells under ischemic stress. Transmission electron microscopy (TEM) imaging reveals distinct apoptotic cleft formation in untreated cardiac cells under ischemic conditions, which is markedly reduced in cells treated with BT (**Figure 6A**). Furthermore, BT treatment effectively downregulates genes associated with early to intermediate apoptosis. it significantly reduces the expression of caspase 8, caspase 9, and caspase 7, with a trending reduction in caspase 3 expression (**Figure 6B-6D, 6F** & **6D**). However, no significant change is observed in caspase 6 expression (**Figure 6E**). The anti-apoptotic effects of BT are further supported by a comparative analysis of the pro-apoptotic protein Bax and the anti-apoptotic protein Bcl-2. BT treatment significantly upregulates Bcl-2 expression, while Bax expression shows a near-significant reduction, collectively contributing to reduced apoptosis in cardiac cells under ischemic stress (**Figure 6G, 6H**). This effect is reinforced by real-time measurements of caspase 3/7 activity, which demonstrate a sustained decrease in caspase 3/7 activity in BT-treated ischemic cardiac cells compared to untreated controls (**Figure 6I**). A comparative profiling of caspase 3/7 activities in cardiac cells treated with BT, palmitate, a lipid mixture, and resveratrol further highlights the superior efficacy of BT as a metabolic therapy (**Figure S6**). Thus, beyond its role in supporting mitochondrial and metabolic recovery, BT shows significant potential in reducing cell apoptosis by effectively lowering mitochondrial ROS levels in cardiac myocytes under ischemic stress.

**Figure 6.**
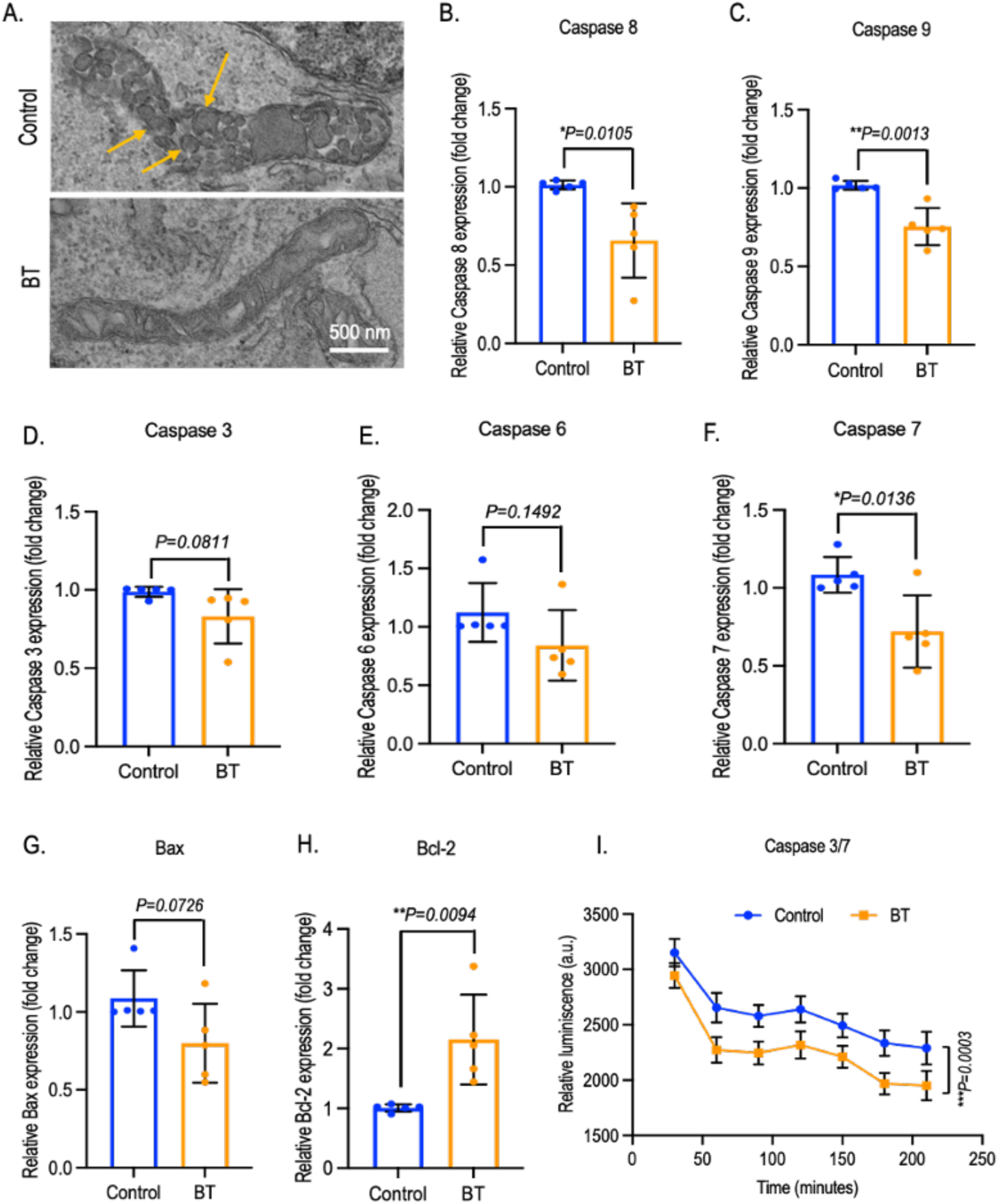
BT limits apoptotic cell death in cardiac myocyte. **(A)** a mitochondrial apoptotic fragmentation in BT and mock-treated perfused cardiac myocytes has been illustrated under electron microscope (Scale bar – 500nm). Significant changes in early apoptotic genes includes-Caspase 8 and Caspase 9 are depicted and quantified by qRT-PCR analysis in figure (**B**) and (**C**) respectively. Changes in intermediate to late apoptotic genes includes-Caspase 3 **(D),** Caspase 6 **(E)** and Caspase 7 (**F)** are depicted quantitively. Significant changes in Bcl-2 family associated apoptotic gene, Bax (**G**) and anti-apoptotic gene, Bcl-2 (**H**) are shown quantitively. A real time measurements of cardiac cell-based Caspase 3/7 **(I)** activities over time are depicted and quantified using bioluminescence assay kit. Data from all BT treatments are quantified and plotted based on results from three to five independent experiments, and comparisons were made against the control group. Each dot in gene expression graph represents an independent experiment, with the mean value derived from three identical replicates. Bar represents the mean ± SD (standard deviation). Statistical analyses are conducted using GraphPad Prism Version 10.3.1 (464), and significance level are denoted as \**p*<0.05, \*\**p*<0.01, based on Student’s t-test. qRT-PCR; Quantitative reverse transcription polymerase chain reaction, Bax; Bcl-2 Associated X-protein, Bcl-2; B-cell lymphoma-2.

## 4. Discussions

Despite a recent increase in annual heart transplants in the United States (around 3,500 last year and a global total of 5,000), the heart transplant waiting list continues to grow rapidly^48^. Shockingly, over 98% of heart-transplant waitlist candidates do not receive transplant each year. However, mechanical circulatory support devices have shown to reduce pretransplant mortality, yet waitlist outcome persist^49^. Additionally, 20% of waitlist candidates die or become too ill for surgery despite surplus donor hearts according to UNOS/OPTN reports^50^. Therefore, efficient allocation of donor hearts could save thousands of lives annually and potentially eliminate the waitlist. Although, most transplanted hearts come from donations after brain death (DBD), yet studies show no significant clinical differences in outcomes compared to DCD hearts^51^. Notably, DCD hearts offer advantages such as shorter wait times, increased transplant rates, and access to younger donors, driving growing interest in their use^52^.

Our findings reveal that under-performing DCD hearts during EVHP exhibit impaired metabolism, marked by uncoupled glycolysis and severe depletion of free fatty acids-leading to rapid ATP loss, and heightened PGD risk^32^. To address these challenges, we optimized a BT-based metabolic therapy in an IVCP model of primary cardiac myocytes, mirroring EVHP protocols to targets I/R-induced mitochondrial and metabolic dysfunction to restore ATP production under ischemic stress and bolster cellular resilience.

During organ transplantation, ischemia disrupts the electron transport chain (ETC), diminishing ATP production, causing electron leakage, and depleting oxygen levels due to inability of alternative energy resource adaptability^53^. This shift to anaerobic metabolism leads to ATP breakdown into ADP and AMP, accumulating inorganic phosphate, which promotes membrane permeabilization, while increasing lactic acid and lowering mitochondrial bioenergetics. Upon reperfusion, oxygen reintroduction generates reactive oxygen species (ROS) that damage proteins, lipids, and DNA, exacerbating mitochondrial dysfunction and triggering apoptosis^54,55^. Herein, cardiac cells efficiently metabolize butyrate, producing acetyl-CoA to fuel the mitochondrial TCA cycle and the ETC complex, thereby sustaining ATP synthesis even under stress conditions. This metabolic adaptation facilitates a shift from glycolysis to mitochondria-driven fatty acid oxidation in stressed cardiac cells, providing a readily available energy source. As a result, it compensates for the metabolic demands typically disrupted by (I/R)-induced mitochondrial injury. This metabolic switch is evidenced by elevated glucose levels, coupled with significant reductions in lactate and alanine production. Besides, BT effectively upregulates α-ketoglutarate levels, which play a cardioprotective role by restoring mitochondrial function, reducing ROS under I/R stress beyond the promotion of cardiac myocyte proliferation and regeneration^41^. Moreover, the quality of mitochondria is essential for sustaining efficient ATP production and preserving optimal cardiac health^56^. In this regard, BT effectively inhibits class 1 histone deacetylase activity and ameliorates mitochondrial biogenesis and oxidative phosphorylation through the upregulation of key proteins including PGC1α, SIRT1, AMPK, NRF1/2 and TFAM in this pathway^57,58^. This, in turn, modulates proteins involved in mitochondrial fission, fusion, and mitophagy, mitigating I/R-induced mitochondrial damage by repairing or eliminating damaged mitochondria that produce excessive ROS. By increasing the expressions of mitophagy-related proteins PINK1 and Parkin, BT enhances mitochondrial quality, recycles damaged mitochondria, reduces ROS, and protects cardiac myocytes from apoptosis. It successfully suppresses early apoptotic pathways, aiding metabolic restoration and preserving energy production during ischemic injury.

In conclusion, BT effectively alleviates mitochondrial damage in cardiac myocytes by enhancing mitochondrial biogenesis, reducing ROS production and preventing cardiac apoptosis. This extends beyond its role in supporting metabolic recovery and ATP production under I/R-induced mitochondrial and metabolic dysfunction. These findings suggest that BT could be a promising metabolic therapy to increase the pool of viable transplant organs by mitigating I/R-induced metabolic dysfunction. However, further research is warranted to gain deeper insights into its safety and efficacy in preclinical trials.

## 5. Limitation, Clinical significance and Future direction

While BT has shown promising outcomes in metabolic therapies during IVCP with human cardiac myocytes, our study is not without limitations. The optimization of BT was conducted exclusively in primary cardiac cells, which, although valuable, may not fully replicate the physiological complexity of the human heart. This highlights the need for further investigations using preclinical models to validate the safety and efficacy of BT in metabolic therapy. Moreover, advancing this research in more sophisticated models is crucial to gain deeper insights into its therapeutic potential.

Nonetheless, our study successfully integrates both fundamental and translational approaches to evaluate and enhance cardiac function. This approach is particularly valuable in addressing metabolic dysfunction in underutilized DCD hearts, offering a pathway to advance metabolic therapies aimed at restoring function in hearts currently deemed non-transplantable. This effort not only aims to enhance metabolic therapy during emergencies but also addresses critical knowledge gaps in managing PGD-associated metabolic dysfunction. Such advancements could lead to the development of more effective strategies for metabolic therapy, ultimately improving organ viability and transplantation outcomes. Building on these successes, we aim to further refine EVHP, transforming it into a more accessible and precise platform for the assessment of donor hearts prior to transplantation. Importantly, the application of metabolic therapies involving short-chain fatty acids holds promise for optimizing mitochondrial metabolic function. These strategies could serve as an influential tool, particularly for researchers focusing on direct mitochondria transplantation as a therapeutic avenue.

## Supporting information

Supplementary details

## Acknowledgement

We thank the Harvard Center for Mass Spectrometry Core facility for their assistance with cell-based metabolomics studies. We also extend our thanks to the HMS Electron Microscopy Core facility for their expertise and assistance in mitochondrial morphology studies.

## Ethics and consent

This study was conducted under protocol number 2022P000405, approved by the Massachusetts General Brigham Institutional Review Board, and was granted exemption from requiring written informed consent. The study adheres to the ethical principles outlined in the Declaration of Helsinki.

## Author Contributions

Conceptualization and experiment design: BR and SAR. Methodology, investigation and visualization: BR, WAM and SRL. Data curation and writing original draft: BR. Writing review and editing: BR, WAM, SRL, SNT, AO and SAR, Fund acquisition and study supervision: SAR. All the authors have read and approved the final manuscript.

## Data availability

All data supporting this study are available upon reasonable request to the authors.

