## Supplementary details for "Improving Cardiac Resilience to Ischemia/Reperfusion: The Role of Butyrate in Mitochondrial and Metabolic Recovery": Suppoting informations.pdf

**Supporting information**
**for**

### Improving Cardiac Resilience to Ischemia/Reperfusion: The Role of Butyrate in Mitochondrial and Metabolic Recovery

Bhuban Ruidas[1], William A. Michaud[1], Sar R. Lindner[2], Asishana A. Osho[1], Shannon N. Tessier[3], Seyed Alireza Rabi[1#]

### Table of Contents


1. **Materials and Methods**………………………………………………………………… **P3-P9**
    *1.1. Ex vivo DCD hearts perfusion*
    *1.2. Primary cell lines and materials*
    *1.3. Cell viability assay*
    *1.4. Perfusate-based electrometabolites and cell-based metabolomics assay*
    *1.5. Mitochondrial morphology analysis*
    *1.6. Cell-based ATP quantification and imaging*
    *1.7. NAD+/NADH analysis*
    *1.8. Mito-respiratory complex I and II quantification*
    *1.9. ADP/ATP analysis*
    *1.10. mRNA extraction, cDNA synthesis and gene expression analysis*
    *1.11. Quantification of hypoxia and mitochondrial ROS*
    *1.12. Caspase activity assay for apoptosis detection*
    *1.13. Immunocytochemistry*
    *1.14. Statistical analysis*

2. **Supplementary Tables**…………………………………………………………. …. **P10-P13**
    *2.1. Table S1: Comparative table of cell-based metabolites*
    *2.2. Table S2: Comparative table of metabolism-associated gene expressions*
    *2.3. Table S3: Mitochondrial ETC complex-associated gene expressions*
    *2.4. Table S4: Mitochondrial DNA synthesis, biogenesis and apoptosis-associated gene expressions*

3. **Supplementary Figures**……………………………………………………… …. **P14-P19**
    *3.1. Figure S1: Comparative cytotoxicity profiles of BT with therapeutic dose comparative and supporting metabolomics profiles*
    *3.2. Figure S2: BT metabolism and glycolysis associated genes expressions*
    *3.3. Figure S3: Mitochondrial ETC complex-associated gene and comparative ADP/ATP ratio profiles*
    *3.4. Figure S4: Mitochondria quantification and mitochondria DNA synthesis associated gene expressions*
    *3.5. Figure S5: Mitochondrial biogenesis associated genes and hypoxia quantification*
    *3.6. Figure S6: A comparative apoptosis profiles with butyrate and different grade of fatty acid and standard control treatment.*

4. **References**……………………………………………………………………… **P20**

#### 1. Materials and methods

##### *1.1. Ex Vivo DCD hearts perfusion*

Our study aligns with the established DCD heart transplantation methodology. The DCD procurement and perfusion process has been detailed previously[1]. Briefly, after obtaining appropriate consent, the donor heart is collected following cessation of life support, a sternotomy, and drainage of donor blood. The heart is then vented, the aorta is cross-clamped, and it is flushed with 1L of 4°C del Nido cardioplegia solution. It is subsequently explanted and placed in the OCS machine for ex vivo perfusion at 34°C. Coronary flow and aortic pressures are continuously monitored, and perfusate samples are collected for experimental analysis using standard protocols. A baseline (BL) sample is taken either pre-OCS placement or within 5 minutes post-placement. The second sample is drawn 1–2 hours after the start of perfusion, and the final sample is collected during precooling and flushing at the recipient's operating room. Our observations indicate that most underutilized hearts exhibit severe depletion of energy and metabolic substrates, including fatty acids, amino acids, and ketones, highlighting the need for immediate therapeutic intervention[2]. To address this, we introduced a metabolic therapy in an *in vitro* cardiomyocyte perfusion (IVCP) model using primary cardiac myocytes extracted from the left ventricles of adult human hearts, aligning with the EVHP methods used for DCD hearts.

##### *1.2. Primary cell lines and materials*

Human cardiac myocytes, HCM and AC10 isolated from adult ventricles were procured from PromoCell (**#12810**) and ATCC (**#CRL-3569**) respectively. HCM were grown in myocyte growth medium (PromoCell C-22070) and AC10 cells were grown in Dulbecco's Modified Eagle Medium/Nutrient Mixture F-12, DMEM-F12 (ATCC 30-2006) with 12.5% FBS (ATCC 30-2020), 1X Insulin-Transferrin-Selenium (ITS -G) and 2% Horse Serum (Gibco Catalog # 16050130). For metabolic therapy experiments, SCFAs, Na-Butyrate (**B5887-250MG**; Sigma-Aldrich), negative inhibitor, Na-palmitate (**P9767-5G**; Sigma-Aldrich) and lipid mixture 1 (**L0288-100ML**; Sigma-Aldrich) and positive control, Resveratrol (**554325-25MG;** Sigma-Aldrich) were used and assessed using standard methods. Lipid Mixture 1 contains non-animal-derived fatty acids (arachidonic acid at 2 µg/mL, and 10 µg/mL each of linoleic, linolenic, myristic, oleic, palmitic, and stearic acids), along with 0.22 mg/mL cholesterol, 2.2 mg/mL Tween-80, 70 µg/mL tocopherol acetate, and 100 mg/mL Pluronic F-68.

##### *1.3. Cell viability assay*

Therapeutic dose of butyrate (BT) was confirmed and quantified varying different concentration using Bio-Rad TC20 automated cell counter (Cat#**1450101)**. Briefly, 2 x$10^4$ cells were seeded and grown to be confluent in 24 well plate at 37ºC with 5% $CO_2$ for 24-48hr of incubation. Confluent cells were then mock treated or treated with different concentrations (0.1, 0.25, 0.5, 0.75, 1, 2, 5 mM respectively) for 1hr. After the treatment, cells were washed twice with 1X PBS and trypsinzed the cells for the detachment from well base. Then trypsinzed cells were stained with trypan blue (dilution; 1:1) and loaded into the cell counter slide to record the cell viability percentage using

Bio-Rad TC20 automated cell counter. Resulting data were normalized and quantified comparing the data of untreated control.

*1.4. Perfusate-based electrometabolites and cell-based metabolomics assay*

To analyze perfusate-derived electrometabolites and cell-based metabolomics, 5x $10^4$ cells were seeded in 12-well plate and grown to confluence before performing IVCP, with or without BT treatment. For electrometabolite analysis, perfusate samples were collected at different stages of IVCP: from baseline hypoxia, during cold ischemia with UW solution, and during warm ischemia with cell media. These samples were then stored at -80°C until electrometabolites profiles were recorded using the Siemen's RAPIDPoint ®500 systems. All the data were quantified relative to control perfusate.

For cell-based metabolomics analysis, cells were trypsinzed and pelleted down to discard the cell lysate. Next the cell pellets were washed with 1X PBS and re-pelleted to store in prechilled methanol (**MX0486; Sigma-Aldrich**: mass spectrometry grade) at -80°C prior to dropping it at core facility of Harvard Center for Mass spectrometry. Briefly, cells were processed with biphasic extraction method for metabolomics analysis in glass vials (VWR 66011-085; 8ml in volume) following the procedure of Bligh and Dyer extraction. Cells were homogenized in 2ml of methanol and transferred to 8ml of glass vials. Next, added 4ml of cold chloroform and 2ml of water and vortexed for 1min followed by an ultrasound bath sonication for 3-5 min to homogenize the cells. Finally, the homogenized cells were centrifuged at 3000 rcf for 10 minutes to separate the aqueous phase, which was then used for metabolomics analysis. Calibration curves with pure standards were prepared for all quantified compounds, and isotope distribution was determined. Samples and calibration curves were prepared by mixing 20 μL of sample with 30 μL of internal standard solution (containing 10 μM D4-Succinate, D11-Methamphetamine, and D11-Amphetamine in acetonitrile) in glass vials. Next, the samples were analyzed using a Q-Exactive Plus Orbitrap MS (Thermo Scientific, Waltham, MA, USA) coupled to an Ultimate 3000 LC system (Thermo Scientific, Waltham, MA, USA). LC separation was performed on a Sequant ZIC-pHILIC column (2.1 × 150 mm, 5 μm, Millipore, Billerica, MA, USA) at a flow rate of 200 μL/min with buffer A (20 mM ammonium carbonate, 0.1% ammonium hydroxide in water) and buffer B (97% acetonitrile in water). The gradient started at 0% to 60% buffer B over 20 minutes, followed by 100% buffer A for 10 minutes. After holding at 100% A for 5 minutes, the gradient was ramped back to 100% buffer B over 5 minutes, followed by 10 minutes of re-equilibration. All samples were run in MS1 mode with polarity switching. MS2 data was subsequently acquired from a representative subset using top 5 data-dependent MS-MS scans in both positive and negative modes on separate runs. The same MS1 data was used for both quantification of the target list and untargeted analysis. For the target list, TraceFinder (Thermo Scientific, Waltham, MA, USA) was used to integrate each target, with quantification based on calibration curves run with pure standards (using the internal standards). Isotope distribution for each target was obtained by integrating the corresponding ion signals. For untargeted analysis, Compound Discoverer 2.0 (Thermo Scientific, Waltham, MA, USA) was used with the same MS1 data set, incorporating MS2 data. Compound identification was based on MS2 matches with the MZcloud library. Compounds without a strong MZcloud match were screened against the KEGG database, using accurate mass only (within 5 ppm), to identify potential candidates and highlight differentially expressed pathways. The MS data file containing identified molecular structures (within 5 ppm) was analyzed using Mummichog software and IPA, as described elsewhere[3].

*1.5. Mitochondrial morphology analysis*

After completion of normothermic IVCP, cells with mock and BT-treated cells were washed twice in 1X PBS and trypsinzed to collect the cell pellets. Next, the pellets were fixed in 2.5% glutaraldehyde/4% paraformaldehyde in 0.1 M Cacodylate buffer and stored in 4°C prior to dropping off at Harvard Medical School (HMS) Electron Microscopy (EM) Core Facility for the imaging of mitochondrial morphology and distribution. Fixed cells were washed in 0.1M cacodylate buffer and postfixed with 1% Osmium Tetroxide (OsO4)/1.5% potassium ferrocyanide (KFeCN6) for five hours, washed in water 3x and once in 50mM of Maleate buffer pH 5.15 (MB), and incubated in 1% uranyl acetate in MB ON at 4°C. Then samples were washed once in MB and 2x in water followed by a subsequent dehydration in a series of graded ethanol (10min each; 50%, 70%, 95%, 2x 10min in 100%). The samples were then put in propylene oxide (PO) f2x over 1hr and infiltrated ON in a 1:1 mixture of PO and TAAB Epon (TAAB Laboratories Equipment Ltd, https://taab.co.uk). The following day, the samples were embedded in TAAB Epon and polymerized at 60°C for 48hr. Next, ultrathin sections (70 nm) were cut on a Reichert Ultracut-S microtome, collected on copper grids and stained with 0.2% lead citrate. Grids were imaged by a Tecnai G2 Spirit BioTWIN TEM equipped with an AMT Nanosprint 43-MKII camera. All chemicals (except the TAAB Epon) are from EMS (***Sciences:www.emsdiasum.com***). Mitochondrial morphology and distribution were quantified using ImageJ software as mentioned elsewhere[4].

*1.6. Cell-based ATP quantification and imaging*

Quantifications of ATP in mock and/or BT-treated perfused-cardiomyocytes were quantified using cell-based ATP-red florescence probe (**Merck; SCT045**) according to the manufacturer protocol. Briefly, 1x $10^3$ cells were seeded in 96 well plate and grown to be confluent prior to perform IVCP with or without BT treatment. Completing IVCP, cells were washed with 1X PBS twice and stained with ATP-red fluorescence probe (5µM in concentration diluting in serum free media) and incubated at 37°C for 15 min. Next, the fluorescence in cells were recorded using microplate spectrophotometer (BioRad) at excitation/emission wavelength of 510nm/570nm.
For ATP imaging, 5x$10^4$ cells were seeded on a coverslip embedded in 12 well plate and grown to be confluent prior to perform IVCP with or without BT treatment. Coverslips with cells were washed with 1X PBS twice and stained with ATP-red fluorescence probe (5µM in concentration diluting in serum free media) and incubated at 37°C for 15 min. After the staining, cover slips with cells were washed with 1X PBS and fixed with DAPI in mounting media (**Abcam; ab104139**) over glass- slide followed by the side shielding with nail-polish and capture the images under EVOS-microscope (ThermoFisher Scientific).

*1.7. NAD+/NADH analysis*

Cellular metabolic state was confirmed measuring intracellular redox regulator NAD+/NADH ratio using NAD/NADH assay kit (**MAK468; Sigma-Aldrich**) following supplier's protocol. Briefly, 2x$10^3$ cells were seeded in 24 well plate and grown to be confluent prior to perform IVCP with or without BT treatment. Completing IVCP, cells were trypsinzed followed by 1X PBS wash and pelleted the cells by centrifugation for 10min at 125 x g at room temperature. Following this, cells were homogenized in a 1.5ml Eppendorf tube with either 100µl of NAD extraction buffer for

NAD determination or 100 µl of NADH extraction buffer for NADH determination at 60°C for 5 min. Next, 20µl of assay buffer and 100µl of the opposite extraction buffer were added to neutralize the extracts and centrifuged the samples at 14000 x g for 5min at room temperature. Next, 40µl of each supernatant were transferred into a separate wells of a clear bottom 96-well plate for NAD/NADH assays and recorded optical density (OD) at 565nm immediately as $OD_0$ and after 15 of incubation as $OD_{15}$. Finally, calculated the NAD and/or NADH concentration of the sample following the formula: NAD/(H) (µM) = $[(\Delta OD_{sample} - \Delta OD_{blank})/ \text{Slope} (\mu M^{-1})]*DF$. Where, $\Delta OD_{sample}$= Change in OD values of sample between zero minutes and 15 minutes, $\Delta OD_{blank}$ = Change in OD values of Blank (Standard #4) between zero minutes and 15 minutes and DF = Sample dilution factor (DF = 1 for undiluted Samples). This colorimetric assay kit included assay buffer (MAK468A), NADH extraction buffer (MAK468B), NADH extraction buffer (MAK468C), Enzyme A (MAK468D), Enzyme B (MAK468E), Lactate (MAK468F), MTT Solution (MAK468G) and NAD standard (MAK468H).

*1.8. Mito-respiratory complex I and II quantification*

Using Mitochondrial Complex I (**Cayman chemicals; 700930**) and Mitochondrial Complex II (**Novus biologicals; NBP3-25854**) activity assay kit, Mito-respiratory Complexes I and II were quantified in mock or BT-treated cardiac myocytes after completion of IVCP following manufacturer protocols. For Mito-respiratory Complex I, 1x $10^3$ cells were seeded in 96 well plate and grown to be confluent prior to perform IVCP with or without BT treatment. Next, cells were washed with 1X PBS followed by the incubation with 100µl assay reagents at 25°C to record the kinetic read at 340nm every 30sec for 15min using microplate reader (BioRad). Finally, determined the % of activity relative to the vehicle control using the equation: Complex I activity (%) = [Rate of Sample wells/Rate of Vehicle Control] *100. Herein, assay reagents were prepared as per supplier's protocol and it includes Complex I activity buffer, mitochondrial inhibitor, FF-BSA assay reagents, bovine heart mitochondria assay reagent, NADH and Ubiquinone assay reagent. Rotenone and antimycin A were used as mitochondrial inhibitor.
For Mito-respiratory Complex II, 1x $10^3$ cells were seeded in 96 well plate and grown to be confluent prior to perform IVCP with or without BT treatment. Next, cells were washed with 1X PBS and collected to analyze Respiratory Complex II as per supplier's protocol (***https://resources.novusbio.com/manual/Manual-NBP3-25854***). Following the sample preparation, blank well and sample well in 96-well plate (provided) were incubated at 37 °C for 3 min with 190µl of reaction working solution to each well. Next, added 20µl of reagent 2 and sample to the blank well and sample well respectively prior to measure the OD value of each well at 600nm with microplate reader and recorded as $A_1$. Second measurement were taken after 3 min of incubation and recorded as $A_2$. Finally quantified the Mitochondrial Complex II activity (U/gprot) using the equation; = $[(\Delta A_{sample} - \Delta A_{blank}) \times V_{total} \times f / (V_{sample} \times 21.8^* \times T \times C_{pr}) \times 1000^*]$. Here, $\Delta A_{sample}$ denotes the change OD value of sample (A1 - A2), $\Delta A_{blank}$ denotes the change OD value of blank (A1 -A2), f denotes dilution factor of sample before test, $V_{total}$ denotes the volume of the reaction system, 0.21 ml, $V_{sample}$ denotes the volume of the sample, 0.02 ml, 21.8* denotes molar absorption coefficient, T denotes time of reaction, $C_{pr}$ denotes the concentration of protein in sample, gprot/L and 1000*:1 mmol/L=1000 µmol/L. Preparation of working solution were followed by suppliers protocol which includes extraction solution A and B, inhibitor, buffer solution and substrate A, B and C. detailed information can be find in above mentioned link of suppliers protocol.

*1.9. ADP/ATP analysis*

An increased levels of ATP and decreased levels of ADP signify proliferating cells. ADP/ATP ratio in primary cardiac myocytes undergoing IVCP with or without BT treatments were investigated using ADP/ATP ratio assay kit (**MAK135; Sigma-Aldrich**) following suppliers' protocol. Briefly, 1x $10^3$ cells were seeded in 96 well plate and grown to be confluent prior to perform IVCP with or without BT treatment. Completing IVCP, cells were washed with 1X PBS twice and incubated with freshly prepared ATP reagents (Assay buffer, Substrate, Co-substrate and ATP enzyme with 95:1:1:1 ratio) for 1min to record luminescence (relative light units) on a luminometer (BioRad) for the ATP assay ($RLU_A$). Next, incubated the plate for additional 10 min in room temperature and again recorded the luminescence for ATP ($RLU_B$). This reading provides the background prior to measure ADP (i.e., the residual ATP signal). Following this, immediately added 5µl of freshly prepared ADP reagent (Water and ADP enzyme with 5:1 ratio) to each well and mixed by tapping the plate. After 1 minute, recorded the luminescence ($RLU_C$) and quantified the ADP/ATP ratio using the formula: ADP/ATP ratio = ($RLU_C$ - $RLU_B$)/ $RLU_A$. In this study. ATP reagents were prepared mixing assay buffer (MAK135A), substrate (MAK135B), co-substrate (MAK135C) and ATP enzyme (MAK135D) with the ratio of 95:1:1:1, whereas ADP reagent were prepared by 1: 5 dilution of ADP enzyme (MAK135E) to water.

*1.10. mRNA extraction, cDNA synthesis and gene expression analysis*

mRNA was extracted from mock- or BT-treated cells following IVCP using an RNA isolation kit (**Zymo Research**) according to the manufacturer's protocols. The mRNA was then quantified and converted into cDNA via polymerase chain reaction (PCR) using a SimpliAMP Thermal Cycler (Applied Biosystems), following standard procedures. Genes associated with glycolysis, mitochondrial dynamics, the ETC complex, apoptosis, and BT metabolism were quantified. Specific forward and reverse primers for all genes were designed and synthesized. Gene expression profiles were analyzed using quantitative reverse transcription polymerase chain reaction (qRT-PCR; Applied Biosystems, Waltham, MA, USA) with SYBR Green (**Thermo-Fisher Scientific, USA**). Gene expression levels were calculated using the ΔΔCT method or the comparative CT method. ΔCT values of the target genes were determined by subtracting the ΔCT values of housekeeping genes, and comparative fold changes were calculated using the standard formula: $2^{-[\Delta CT\ (treated)\ -\ \Delta CT\ (calibrated)]}$. Glyceraldehyde 3-phosphate dehydrogenase (GAPDH) and *β*-actin were used as house-keeping genes to normalize the gene expressions. All the primers were obtained from IDT technologies (Iowa, USA). A comprehensive list of primers with level of significance summary is provided in the supporting tables.

*1.11. Quantification of hypoxia and mitochondrial ROS*

Hypoxia in cardiac myocytes undergoing IVCP with or without BT treatment were quantified using Image-iT™ green hypoxia reagent (**ThermoFisher Scientific, #I14834**), while mitochondrial ROS were measured using MitoSOX mitochondrial superoxide indicator (**ThermoFisher Scientific, #M36006**), followed by the counter staining with mitochondria labelling using MitoTracker™ dyes (**ThermoFisher Scientific, #M7512**). All the experiments were carried out following supplier's protocol. Briefly, Briefly, 1x $10^3$ cells were seeded in 96 well plate for fluorescence measurements and 2x $10^3$ cells were seeded in 8 well chamber (μ-Slide 8 well high, ibidi; #80806) grown to be confluent prior to perform IVCP with or without BT treatment. To measure cell hypoxia, cells were washed with 1X PBS twice and incubated with 1μM of hypoxia reagents (freshly prepared in protein free cell media) at 37°C for 30 min prior to capture the image and fluorescence intensities with excitation and emission wavelength at 488nm and 520nm respectively. For mitochondrial ROS measurement, post treatment IVCP cells were washed with warm buffer (HBBS with calcium and magnesium or suitable buffer) thrice and incubated with 1μM of MitoSOX™ green reagents (freshly prepared) for 30 min at 37°C, protecting from light followed by counter staining with MitoTracker™ to assess mitochondrial membrane potential as a factor in mitochondrial accumulation of MitoSOX™ sensors for imaging or without the counter staining for fluorometric analysis at excitation and emission wavelength of 488nm and 510nm respectively. Hanks balanced salt solution (HBSS) with calcium, magnesium were procured from Thermos Fisher Scientific (#14025092).

*1.12. Caspase activity assay for apoptosis detection*

Apoptotic cell death in mock or BT-treated cardiac myocytes undergoing IVCP were investigated and quantified using Caspase-Glo 3/7 assay (**Promega, #G8091-10ml**) systems following suppliers' instructions. Briefly, 1x $10^3$ cells were seeded in white-walled 96 well plate and grown to be confluent prior to perform IVCP with or without BT treatment. Completing IVCP, cells were washed with 1X PBS twice and incubated with 100μl of cell media and freshly prepared caspase-Glo 3/7 reagent (1:1) for 30 min to 3 hours at room temperature after sealing the plate with gentle shaking using a plate shaker at 300-500rpm for 30 sec. Finally, recorded the luminescence using luminometer (BioRad) each 30 min of interval up to 3hr. Caspase-Glo 3/7 reagents were prepared mixing caspase-Glo 3/7 buffer with lyophilized caspase-Glo 3/7 substrate as per manufacturers instruction[5].

*1.13. Immunocytochemistry*

IVCP cells, treated with or without BT, were gently rinsed with pre-warmed PBS and fixed in 4 % paraformaldehyde (**Sigma-Aldrich, St Louis, MO, USA**) for 10-20 min at room temperature. Permeabilization was performed using PBS containing 0.05–0.1% Triton X-100 (PBST; Sigma-Aldrich, St. Louis, MO, USA). To block nonspecific binding, the cells were incubated for 30 minutes in 2% Bovine Serum Albumin (**BSA; Jackson ImmunoResearch**) or 10 % goat serum in PBST. For primary antibody dilution, PBS with 2 % goat serum or 0.5% BSA was used. The following primary antibodies were used at room temperature for 2 hours: rabbit anti-Mitofusin-2 (**Millipore Sigma-M6319, 1:100**), mouse anti-PARK2/Parkin (**Proteintech-66674-1-Ig; 1:100**). After incubation, the primary antibody solution was washed off with PBST three times. Following this, secondary antibody staining was performed in PBST for 1h at room temperature, using Alexa Fluor-488 conjugated secondary antibodies (**Invitrogen, Carlsbad, CA, USA**). For counter staining, nuclei were stained with Hoechst (**10 mg/ml, Invitrogen, Carlsbad, CA, USA**) 1:5000. Imaging was carried out using an EVOS-microscope (ThermoFisher Scientific) and the resulting images were quantified using ImageJ software.

*1.14. Statistical analysis*

All experiments were performed with a n = 3-7 independent replicates. Data are presented as the mean ± standard error of mean (s.e.m) and standard deviation (SD). MS results were correlated standard calibration samples and only results with high confidence level of >90% were included in the analysis. Statistical significance was analyzed using as unpaired two-tailed Student's t-test in Prism (GraphPad Software, La Jolla California USA). Mitochondrial distributions are quantified by Pearson's linear correlation (Two-tailed) analyses the 95% of confidence interval in **figure 4D**. Statistical significance thresholds were set at $p<0.05$ (*), $p<0.01$ (**), $p<0.001$(***), and $p<0.0001$(****).

#### 2. Tables

**Table S1:** Comparative table of cell-based metabolites in perfused cardiac cell with or without BT treatment are depicted and quantified with statistical significance.

| **Metabolites (cell based)** | **Mean spectral area (Control)** | **Mean spectral area (BT)** | **P- value** |
|---|---|---|---|
| Glucose | 6000000 | 9000000 | ***P=0.0025*** |
| Fructose 1,6 bisphosphate | 5e+007 | 3e+007 | ***P=0.4695*** |
| Phosphoenolpyruvate | 200000 | 400000 | ***P=0.6985*** |
| Pyruvate | 2e+008 | 2e+008 | ***P=0.8408*** |
| Lactate | 1e+007 | 6000000 | ***P=0.0111*** |
| Citrate | 1.73E+08 | 1.4E+08 | ***P=0.4971*** |
| Aconitate | 3e+007 | 2e+007 | ***P=0.2933*** |
| α-ketoglutarate | 7E+06 | 2E+07 | ***P=0.1206*** |
| Succinate | 4E+08 | 3E+08 | ***P=0.1901*** |
| Fumarate | 1E+08 | 9E+07 | ***P=0.0666*** |
| Malate | 1E+09 | 6E+08 | ***P=0.6176*** |
| AA-Alanine | 8e+007 | 7e+007 | ***P=0.6649*** |
| AA-Ornithine | 4E+06 | 2E+06 | ***P=0.6707*** |
| AA-Isoleucine + Leucine | 7E+07 | 6E+07 | ***P=0.4226*** |
| AA-Valine | 7E+09 | 7E+09 | ***P>0.9999*** |
| AA-Histidine | 3E+06 | 2E+06 | ***P>0.9999*** |
| AA-Lysine | 1E+07 | 8E+06 | ***P=0.4226*** |
| AA-Glutamate | 5E+06 | 2E+06 | ***P=0.0955*** |
| AA-Glutamine | 2E+06 | 1E+06 | ***P=0.2929*** |
| AA-Proline | 1.3E+09 | 6.25E+08 | ***P=0.1918*** |
| Phosphocholine | 4e+010 | 3e+010 | ***P=0.3107*** |
| Pantothenic acid | 1e+008 | 2e+008 | ***P=0.0434*** |
| Choline | 3e+008 | 4e+008 | ***P=0.0335*** |
| AMP | 3E+08 | 1E+08 | ***P=0.0125*** |
| ADP | 2E+08 | 1E+08 | ***P=0.2605*** |
| ATP | 4E+07 | 1E+08 | ***P=0.1158*** |

**Table S2:** Comparative table of metabolism-associated gene expressions in perfused cardiac cell with or without BT treatment are depicted and quantified with statistical significance.

| **Gene** | **Forward primer (5'-3')** | **Reverse primer (5'-3')** | **Mean Result (Control vs BT)** | ***P*-value** |
|---|---|---|---|---|
| FAT/CD36 | CAGGACGCTGAGGACAACACAGTC | GTACAGATGCAGCCTCATTTCCACC | 1.053 vs 1.239 | ***P*=0.6093** |
| MCT1 | GAGCAATTTCCAGGTATTGAACCATGG | CCTGCTGCAGTAGATACCATTGTTGC | 1.008 vs 1.773 | ***P*=0.1162** |
| CPT1A | GCCTTTCAGTTCACGGTCACTCC | GACTCTGGAAACGGCCAACTGC | 1.055 vs 1.891 | ***P*=0.0523** |
| CPT2 | GGAGATACCTCAGTGCACAGAAGCC | GCTCGAGACTCCGTTGTTCTGAAC | 1.744 vs 3.627 | ***P*=0.1867** |
| ACSM3 | GGAAGATGCTACGTCATGCCAAGTG | GGAGTTGATGAAACATGCCAGTGACAG | 1.132 vs 3.326 | ***P*=0.0479** |
| SCAD1 | GAAGGAGTTGTTTCCCATTGCAGCC | CTGGATCACCAATGCCTGGGAG | 1.026 vs 2.975 | ***P*=0.0077** |
| ECHS1 | CAATGCACTTTGCGATGGCCTG | CCTCACCCAGGTCAAGAAGCCAGTC | 1.041 vs 2.186 | ***P*=0.0757** |
| HADH | GGTAGACCAGACAGAGGACATCCTG | GCTGGACAAGTTTGCTGCTGAAC | 1.018 vs 1.452 | ***P*=0.0952** |
| ACAT1 | CTTTCCTTGCTGCCAGCCAC | GGCCTCTCAAAGTCTTATGTGTGGAC | 1.016 vs 3.708 | ***P*=0.0156** |
| PDH | GGAACGTCTGTTGAGAGAGCGG | CCTGATGGAGCTGCAGACTTACCG | 1.219 vs 1.163 | ***P*=0.8767** |
| PDK4 | CCATGAAGCAGCTACTGGACTTTGG | GTCCCTACAATGGCACAAGGAATCATAG | 1.027 vs 1.737 | ***P*=0.0123** |
| LDHA | GGCAGATGAACTTGCTCTTGTTGATGTC | GCAATCTGGATTCAGCCCGATTC | 1.098 vs 0.896 | ***P*=0.2155** |
| ALAT | GAAGAAGCCTTTCACCGAGGTC | GACGAGGTGTACCAGGACAACG | 1.111 vs 1.115 | ***P*=0.9848** |
| GAPDH | AAGAAGGTGGTGAAGCAGG | GTCAAAGGTGGAGGAGTGG | House-keeping gene | |
| β-Actin | CACCATTGGCAATGAGCGGTTC | AGGTCTTTGCGGATGTCCACGT | House-keeping gene | |

**Table S3:** Mitochondrial ETC complex-associated gene expressions in perfused cardiac cell with or without BT treatment are depicted and quantified with statistical significance.

| Gene | Forward primer, 5'-3' | Reverse primer, 5'-3' | Mean Result (Control vs BT) | *P*-value |
|---|---|---|---|---|
| **NADH dehydrogenase (Complex-I)** | | | | |
| NDUFA1 | CTTCCGGCTGCTTATTTGAATCC | CTGGAGTCTGATGGAAAGAGATAGGC | 1.039 vs 1.641 | ***P*=0.0634** |
| NDUFA4 | CAGAGCCCTGGAACAAACTGGG | GACCTTCATTCTAAAGCAGCG | 1.067 vs 2.035 | ***P*=0.1150** |
| NDUFB3 | GCTCCGGTTGCAGAGTTGAGTGTC | GGAGCTGAATATTACCTGGAGTCCCTG | 1.011 vs 1.611 | ***P*=0.2654** |
| NDUFB5 | GCTGAACTAGCAGAAATTCCAGAAGGC | GAGGAGTTTGGCTTCCAGTCCC | 0.914 vs 1.204 | ***P*=0.1828** |
| NDUFC1 | CACCGCAGTGGAGACTTGTTAACAG | GGCCATATAATGCAGAGTTGACGG | 1.154 vs 1.653 | ***P*=0.2579** |
| NDUFS2 | ACCCAAGCAAAGAAACAGCC | AATGAGCTTCTCAGTGCCTC | 1.021 vs 1.564 | ***P*=0.0499** |
| MT-ND1 | CTACTACAACCCTTCGCTGAC | GGATTGAGTAAACGGCTAGGC | 0.947 vs 1.662 | ***P*=0.0031** |
| MT-ND2 | CATATACCAAATCTCTCCCTC | GTGCGAGATAGTAGTAGGGTC | 0.972 vs 1.484 | ***P*=0.0047** |
| MT-ND3 | TTACGAGTGCGGCTTC | CCTAGTTTTAAGAGTACTGCG | 0.984 vs 1.801 | ***P*=0.0875** |
| MT-ND4 | CCACCTTGGCTATCATCACCC | CTCCACTTATGACTCCCTAAAGCCC | 1.022 vs 2.379 | ***P*=0.0091** |
| MT-ND4L | TAGTATATCGCTCACACCTC | GTAGTCTAGGCCATATGTG | 1.090 vs 2.038 | ***P*=0.0138** |
| MT-ND5 | TCGAATAATTCTTCTCACCC | TAGTAATGAGAAATCCTGCG | 1.079 vs 1.948 | ***P*=0.0459** |
| MT-ND6 | GTAGGATTGGTGCTGTGG | GGATCCTCCCGAATCAAC | 1.016 vs 1.705 | ***P*<0.0001** |
| **Succinate dehydrogenase (Complex-II)** | | | | |
| SDHA | CTGAGGCAGGGTTTAATACAGCATGTG | GTGGCATTTCTACGACACCGTG | 0.961 vs 3.146 | ***P*=0.0388** |
| SDHB | GGCTGGAGACAAACCTCATATGC | CTAGCTTGCACCCGAAGGATTG | 1.028 vs 1.841 | ***P*=0.0548** |
| SDHD | GTTGCTTCGAACTCCAGTGGTC | CTTCCGGCTGCTTATTTGAATCC | 1.001 vs 2.801 | ***P*=0.0232** |
| **Cytochrome C reductase (Complex-III)** | | | | |
| UQCRC1 | AATGGGGCAGGCTACTTTTT | GGTCAAGTCTGCACGAGACA | 1.022 vs 1.982 | ***P*=0.0719** |
| UQCRH | ACTGGAGGACGAGCAAAAGA | TGATGCCCAGATGATGAAGA | 1.040 vs 1.980 | ***P*=0.1305** |
| UQCRQ | AGTGCAGTGGTGTGATCTCG | CTGTGCCCATTTCCTCATCT | 1.419 vs 3.046 | ***P*=0.2868** |
| CYC1 | CCAAAACCATACCCCAACAG | TATGCCAGCTTCCGACTCTT | 1.018 vs 2.692 | ***P*=0.0003** |
| CYTB | TGAAACTTCGGCTCACTCCT | AATGTATGGGATGGCGGATA | 1.051 vs 1.679 | ***P*=0.0016** |
| **Cytochrome C oxidase (Complex-IV)** | | | | |
| COX1 | CTGCTATAGTGGAGGCCGGA | GGGTAGGAGTAGTTCCCTGC | 1.025 vs 1.675 | ***P*=0.0139** |
| COX2 | GCACATGCAGCGCAAGTAGGTCTAC | GGCCACCAATGGTACTGAACCTACG | 1.010 vs 1.788 | ***P*=0.0345** |
| COX3 | CCAATGATGGCGCGATG | CTTTTTGGACAGGTGGTGTGT | 1.038 vs 1.535 | ***P*=0.0131** |
| COX5A | GTCACCTGACCAGAGACAAGG | AGGTCACCGCAAGGACAC | 0.797 vs 1.386 | ***P*=0.1870** |
| COX7A1 | TAAATACCGTTTTACTCCCAAA | CTCGGATTCGTCCACCAC | 1.021 vs 1.479 | ***P*=0.1797** |
| COX7B | AAGGGATTGCAATTACTATAGGTTT | CGTAAAAGGAAAGCACACGA | 1.043 vs 2.208 | ***P*=0.1341** |
| **ATP synthase (Complex-V)** | | | | |
| MT-ATP6 | CCTCCCTCACCAAAGCCCAT | GGTCATGGGCTGGGTTTTACTA | 1.037 vs 2.942 | ***P*=0.0204** |
| MT-ATP8 | CCCCTCTAGAGCCCACTGTAA | GTGGGGATCAATAGAGGGGGA | 1.029 vs 1.721 | ***P*=0.0010** |
| ATP5F1A | GTTCCAGTTGGTGAGGAGCTGTTG | CAGTACCTGGCTCCTTACTCTGGCTG | 1.008 vs 3.725 | ***P*=0.0249** |
| ATP5F1D | CAGATGTCCTTCACCTTCGCCTCTC | CAAACTTGGAGAAGGCCCAGGC | 1.081 vs 2.416 | ***P*=0.0435** |
| ATP5F1E | CAGGTTAGGAGCCCTTGGAGTCC | CTGGGTAGCTGGCAGGATGACTG | 1.010 vs 1.360 | ***P*=0.0122** |
| GAPDH | AAGAAGGTGGTGAAGCAGG | GTCAAAGGTGGAGGAGTGG | House-keeping gene | |
| β-Actin | CACCATTGGCAATGAGCGGTTC | AGGTCTTTGCGGATGTCCACGT | House-keeping gene | |

**Table S4:** Mitochondrial DNA synthesis, biogenesis and apoptosis-associated gene expressions in perfused cardiac cell with or without BT treatment are depicted and quantified with statistical significance.

| Gene | Forward primer, 5'-3' | Reverse primer, 5'-3' | Mean Result (Control vs BT) | *P*-value |
|---|---|---|---|---|
| **Mitochondrial DNA synthesis-associated genes** | | | | |
| HDAC1 | GCTGAAGACCTGGAGACAGATGGC | CTCCTGAGCCAGTGTGAGTGTCTCC | 1.149 vs 0.794 | ***P*=0.0155** |
| POLG 1 | CAAAGCCAGGTGTTCTGACTCCC | CGAGCAAATCTTCGGGCAAG | 0.953 vs 1.510 | ***P*=0.0219** |
| POLG2 | CAGGTGCCTGTTGTCTGGGTTTG | CAAACCAGGCCCTTTGCTACCC | 1.015 vs 1.725 | ***P*=0.0507** |
| TFAM | CACATTTTCCACCTGGTGAT | CACTCCGCCCTATAAGCATC | 1.021 vs 1.395 | ***P*=0.0523** |
| MTIF2 | GAGCATTAAGACAGTGGAGGCATGG | GAGGCATCACTCAGCACATTGGTG | 1.060 vs 1.890 | ***P=0.0144*** |
| **Mitochondrial Biogenesis and mitophagy system-associated genes** | | | | |
| PGC1α | TCAGTCCTCACTGGTGGACA | TGCTTCGTCGTCAAAAACAG | 1.046 vs 1.995 | ***P*=0.0327** |
| AMPK | TTGAAACCTGAAAATGTCCTGCT | GGTGAGCCACAACTTGTTCTT | 1.008 vs 1.454 | ***P*=0.0026** |
| SIRT1 | CGATTGGGTACCGAGATAACCTTCTG | CATGGTTCCTTTGCAACAGCATC | 1.046 vs 1.761 | ***P*=0.0349** |
| NRF1 | GGTGCAGCACCTTTGGAGAA | CCAGAGCAGACTCCAGGTCTTC | 1.026 vs 1.691 | ***P*=0.0308** |
| NRF2 | CCAGGCCATAGACATCAATGAACC | CCCACCATGCTGAATCAGAAGC | 1.073 vs 3.078 | ***P*=0.0993** |
| DRP1 | CAAAGCAGTTTGCCTGTGGA | TCTTGGAGGACTATGGCAGC | 1.007 vs 1.621 | ***P=0.0094*** |
| FIS1 | CCAGGTAGAAGACGTAATCCC | GTCCAAGAGCACGCAGTTT | 0.998 vs 1.552 | ***P*=0.1176** |
| MFN1 | ATGTAACGGACGCCAATC | ATCTTTAGCTTCTACTCCCACT | 1.001 vs 1.547 | ***P*=0.0053** |
| MFN2 | TGCAGGTGTAAGGGACGATT | GAGGCTCTGCAAATGGGATG | 0.995 vs 1.569 | ***P*=0.0008** |
| OPA1 | TGTCCTCCGCAAAGTCAT | TGCTTGGGAGACCCTACA | 1.002 vs 1.640 | ***P*=0.1103** |
| Parkin | GAAGAACCACCTCAGCCTCAGGAG | GCCAGCAATTGTAACCCACACTCTG | 1.011 vs 1.548 | ***P*=0.0003** |
| PINK1 | CATTGGTAAGGGCTGCAGTGCTGC | CCATCTTGAACACAATGAGCCAGGAGC | 1.046 vs 1.979 | ***P*=0.0382** |
| HIF-1α | CCAGATCTCGGCGAAGTAAAGAATC | CTTGGTGCTGATTTGTGAACCC | 1.053 vs 0.854 | ***P*=0.2230** |
| **Apoptosis-associated genes** | | | | |
| Caspase 3 | GGCAATGCTGACCTTCATACCTAGTG | GCACTGGTGTTTGGGCCTAAAC | 0.988 vs 0.831 | ***P*=0.0811** |
| Caspase 6 | GCACCTGCGCAGATAGAGACAATC | GACAAGTGTCACAGCCTGGTTGG | 1.112 vs 0.841 | ***P*=0.1492** |
| Caspase 7 | CAGTGGATGCTAAGCCAGACCG | CTGCATCCTCTTAAGCCATGGAGAAG | 1.084 vs 0.720 | ***P*=0.0136** |
| Caspase 8 | CTGGCAGTGTCCAAGCAACAAGC | GTTTGGTTCCTGGAGTCAGCACCTG | 1.012 vs 0.656 | ***P*=0.0105** |
| Caspase 9 | GGAGTCAGGCTCTTCCTTTGTTCATC | GCTGCATTTCATGGTGGAGGTG | 1.018 vs 0.754 | ***P*=0.0013** |
| Bax | CTGACGGCAACTTCAACTGGG | CATGGGCTGGACATTGGACTTC | 1.086 vs 0.799 | ***P*=0.0726** |
| Bcl-2 | GGACTTCTGCGAATACCGGACTG | CAGTGTGTACAGGGAAACGCAC | 1.006 vs 2.151 | ***P*=0.0094** |
| GAPDH | AAGAAGGTGGTGAAGCAGG | GTCAAAGGTGGAGGAGTGG | House-keeping gene | |
| β-Actin | CACCATTGGCAATGAGCGGTTC | AGGTCTTTGCGGATGTCCACGT | House-keeping gene | |

#### 3. Figures

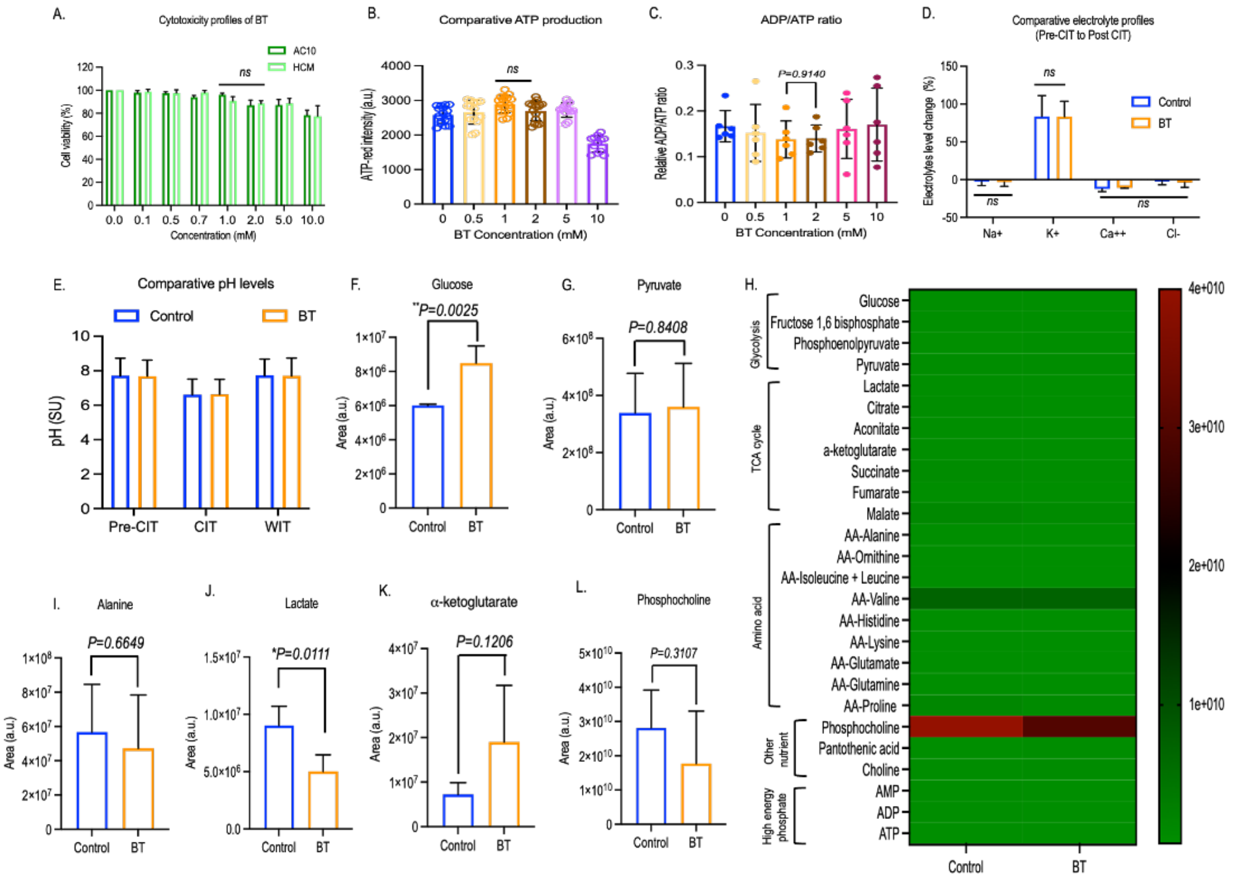


**Figure S1.** Comparative cytotoxicity profiles of BT in IVCP model of primary cardiac myocytes include-HCM and AC10 isolated from human adult ventricles are shown (**A**) followed by the quantification of optimum therapeutic dose of BT through comparative ATP production (**B**) and ADP/ATP ratio analysis (**C**). Cell perfusate-based electro-metabolites profiles include- comparative electrolyte level (**D**) and pH **(E).** Cardiac myocyte-based spectrometric metabolome profiles include- Glucose **(F)**, pyruvate **(G)**, heat map of major cell-based metabolites **(H)**, alanine **(I),** lactate **(J)**, α-ketoglutarate (**K**) and phosphocholine **(L)** are depicted and quantified. All the BT treatment data resulted and plotted based on three to four independent experiments and compared to control. The bar represents the mean ± SD (standard deviation). Statistical analyses were conducted using GraphPad Prism Version 10.3.1 (464), and significance level are denoted as *$p$<0.05, **$p$<0.01 based on Student's t-test. 'ns' denotes no significance. IVCP; In vitro cell perfusion, HCM; human cardiac myocytes.

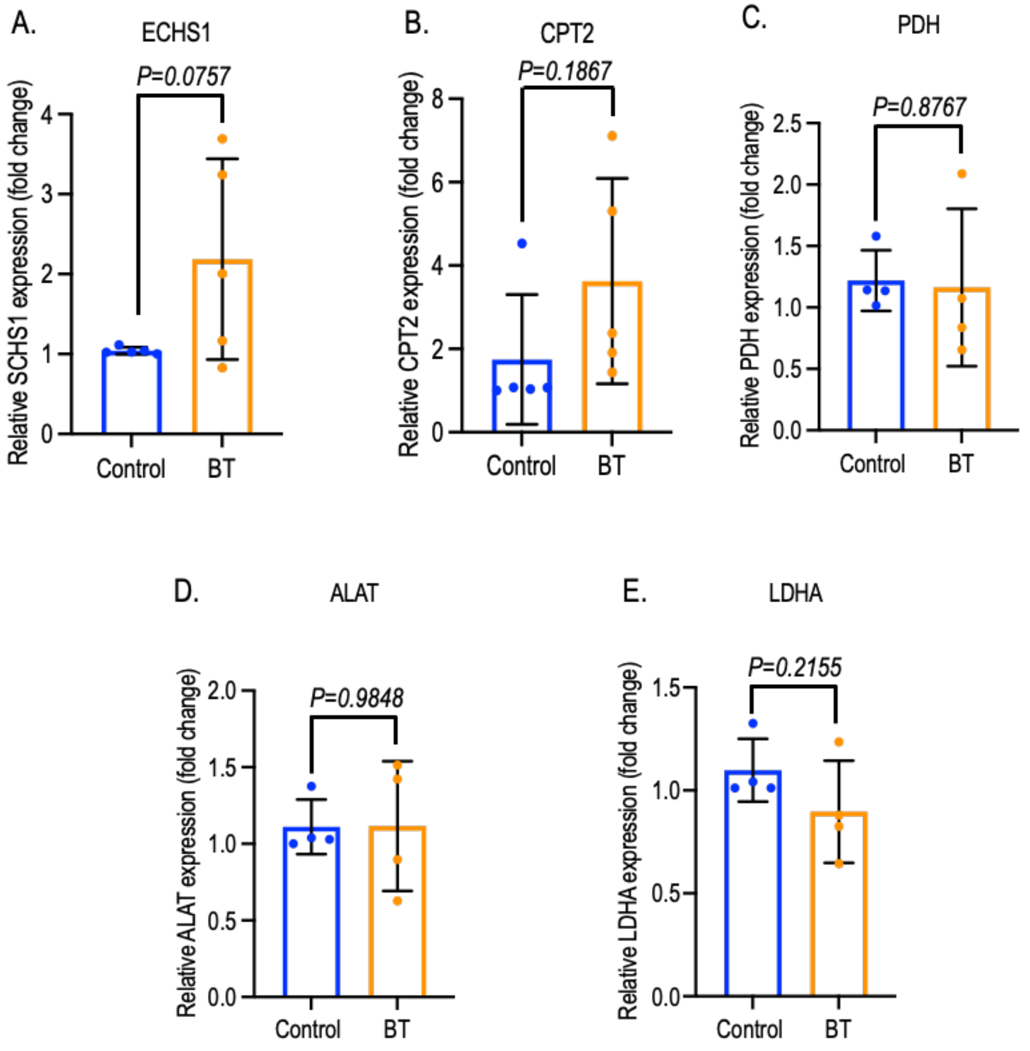


**Figure S2.** In perfused cardiac cells, qRT-PCR based expressions of butyrate (BT) metabolism-associated enzyme, ECHS1 (**A**) responsible for conversion of crotonyl-CoA to 3-hydroxybutyryl-CoA are depicted. (**B**) Expression of inner mitochondrial fat metabolism-associated protein, CPT2 are depicted and quantified. Pyruvate to acetyl-CoA and pyruvate to Alanine includes- PDH **(C)** and ALAT **(D)** are depicted respectively. (**E**) Pyruvate to lactate conversion initiator gene LDHA are shown. All the BT treatment data were quantified and plotted based on three to five independent experiments and compared to control. Each dot in gene expression graph represents an independent experiment, with the mean value derived from three identical replicates. The bar represents the mean ± SD (standard deviation). Statistical analyses were conducted using GraphPad Prism Version 10.3.1 (464), and significance level are denoted as *$p<0.05$, based on Student's t-test. qRT-PCR; Quantitative reverse transcription polymerase chain reaction, PDH; Pyruvate dehydrogenase, LDHA; lactate dehydrogenase A, ALAT; Alanine aminotransferase, ECHS1; Short-chain enoyl-CoA hydratase, HADH; 3-Hydroxyacyl-CoA dehydrogenase.

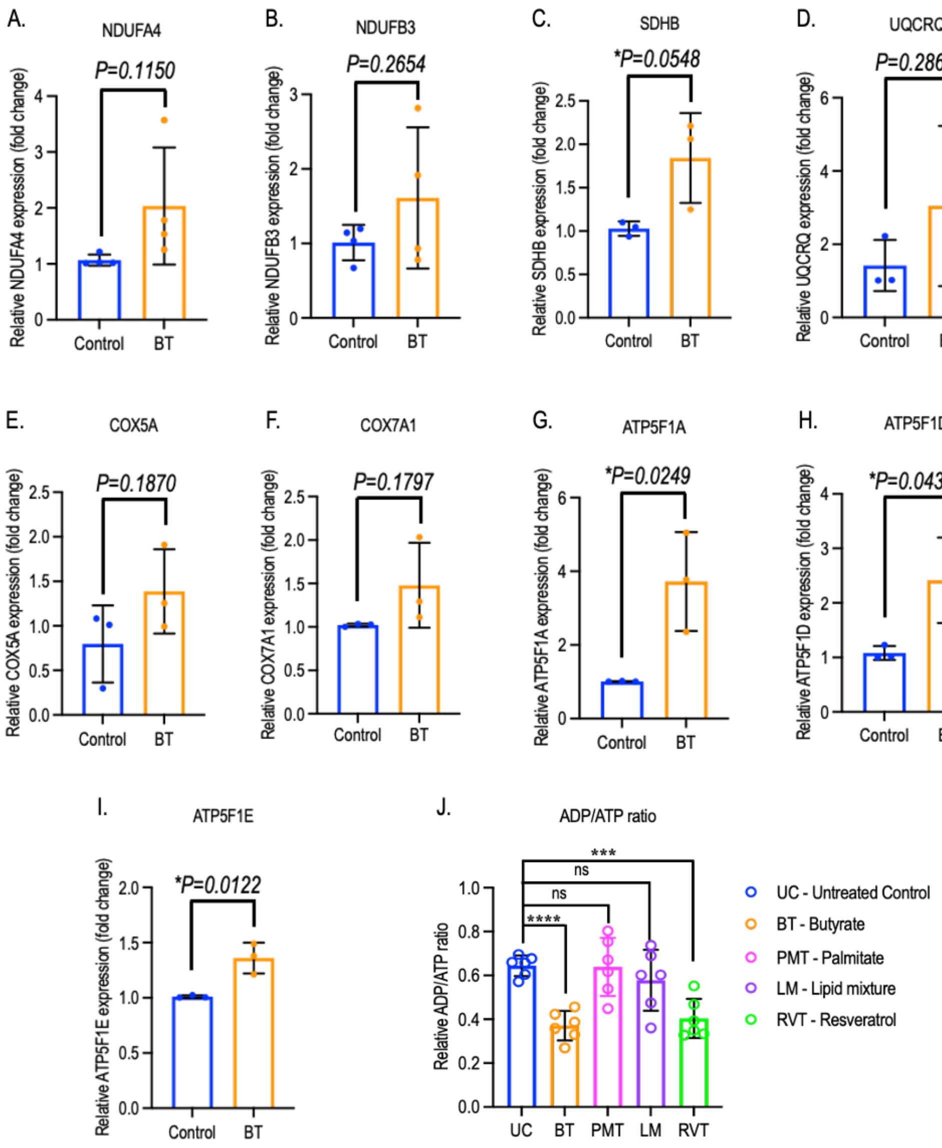


**Figure S3.** In mock or BT-treated perfused cardiac cells, qRT-PCR based expressions of mitochondrial ETC complex-I-associated gene include- NDUFA4 **(A)**, NDUFB3 **(B)** and ETC complex-II-associated gene include- SDHB **(C)** are depicted and qunatified. Expression of other genes**-** UQCRQ **(D)** associated with ETC complex-III, COX5A **(E)** and COX7A1 **(F)** associated with complex-IV and ATP51A (**G**), ATP5F1D (**H**), and ATP5F1E (**I**) associated with complex-V are depicted respectively. (**J**) A comparative diagram of ADP/ATP ratio treated with butyrate and different grade of fatty acid include- long chain fatty acid (palmitate), lipid mixture (0.1%) and standard control, resveratrol are depicted and quantified compared to control. All the BT treatment data were quantified and plotted based on three to five independent experiments and compared to control. Each dot in gene expression graph represents an independent experiment, with the mean value derived from three identical replicates. The bar represents the mean ± SD (standard deviation). Statistical analyses were conducted using GraphPad Prism Version 10.3.1 (464), and significance level are denoted as **p*<0.05, ****p*<0.001, *****p*<0.0001 based on Student's t-test. 'ns' denotes no significance. qRT-PCR; Quantitative reverse transcription polymerase chain reaction, NDUFA4; NADH dehydrogenase (ubiquinone) 1 alpha subcomplex 4, NDUFB3; NADH dehydrogenase (ubiquinone) 1 beta subcomplex 3, SDHB; Succinate dehydrogenase [ubiquinone] iron-sulfur subunit B, UQCRQ; Cytochrome b-c1 complex subunit 8, COX5A and COX7A1; Cytochrome oxidase subunit 5A and 7A1, ATP5F1(A, D and E); ATP synthase F1 (subunit alpha, delta, and epsilon)

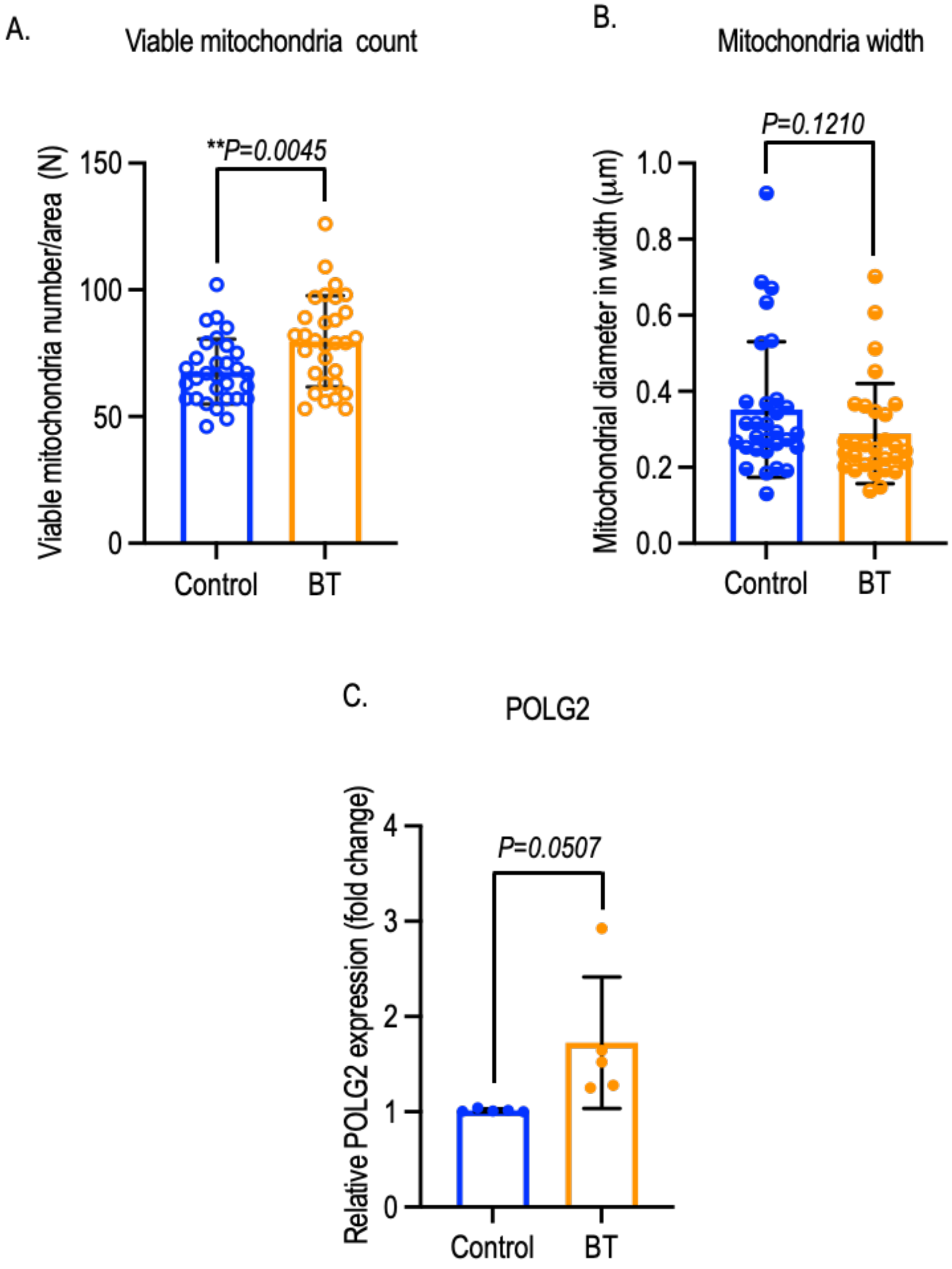


**Figure S4.** (**A**) Viable mitochondria count per area are shown in carding myocytes with or without BT treatment following IVCP methods. (**B**) Quantification of comparative mitochondria width are depicted. qRT-PCR based expressions of mitochondrial DNA replication gene, POLG2 **(C)** for DNA synthesis is depicted quantitively. All the BT treatment data were quantified and plotted based on three to five independent experiments and compared to control. Each dot in gene expression graph represents an independent experiment, with the mean value derived from three identical replicates. The bar represents the mean ± SD (standard deviation). Statistical analyses were conducted using GraphPad Prism Version 10.3.1 (464), and significance level are denoted as *$p<0.05$, **$p<0.01$, based on Student's t-test. IVCP; In vitro cell perfusion, qRT-PCR; Quantitative reverse transcription polymerase chain reaction, POLG2; polymerase gamma 2.

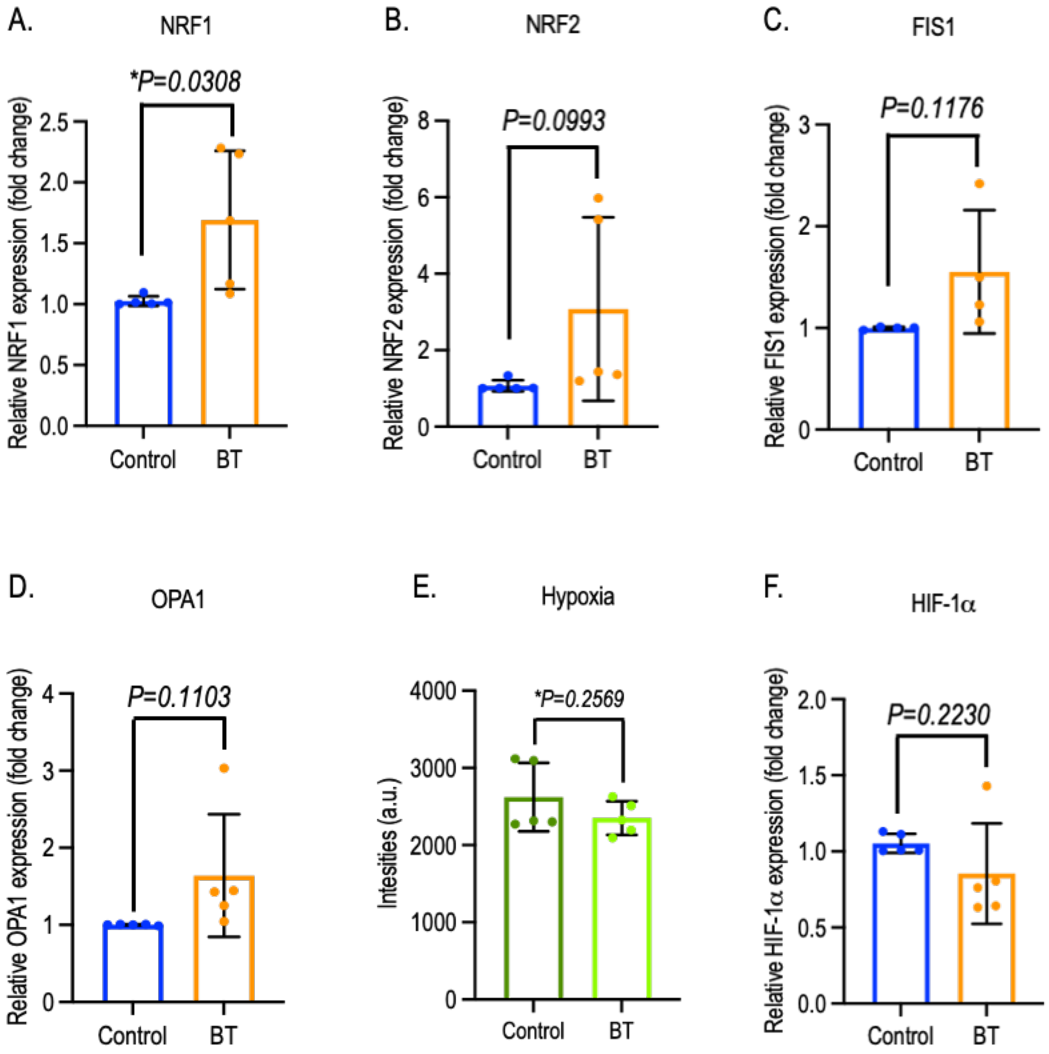


**Figure S5.** In mock or BT-treated perfused cardiac cells, qRT-PCR based expressions of mitochondrial biogenesis associated genes include- NRF1 (A) and NRF2 (B) are depicted. Expression of mitochondrial fission associated gene, FIS1 (C) and fusion associated gene, OPA1 (D) are depicted and quantified. Real time measurement of hypoxia (E) and expression of it's in associative gene, HIF-1α in cardiac myocytes undergoing ischemia/reperfusion are shown and quantified. All the BT treatment data were quantified and plotted based on three to five independent experiments and compared to control. Each dot in gene expression graph represents an independent experiment, with the mean value derived from three identical replicates. The bar represents the mean ± SD (standard deviation). Statistical analyses were conducted using GraphPad Prism Version 10.3.1 (464), and significance level are denoted as *$p<0.05$, **$p<0.01$, based on Student's t-test. qRT-PCR; Quantitative reverse transcription polymerase chain reaction, NRF1; Nuclear Respiratory Factor 1, NRF2; Nuclear factor erythroid 2-related factor 2, FIS1; mitochondrial fission protein 1, OPA1; Optic atrophy 1, HIF-1α; Hypoxia inducible factor 1 alpha.

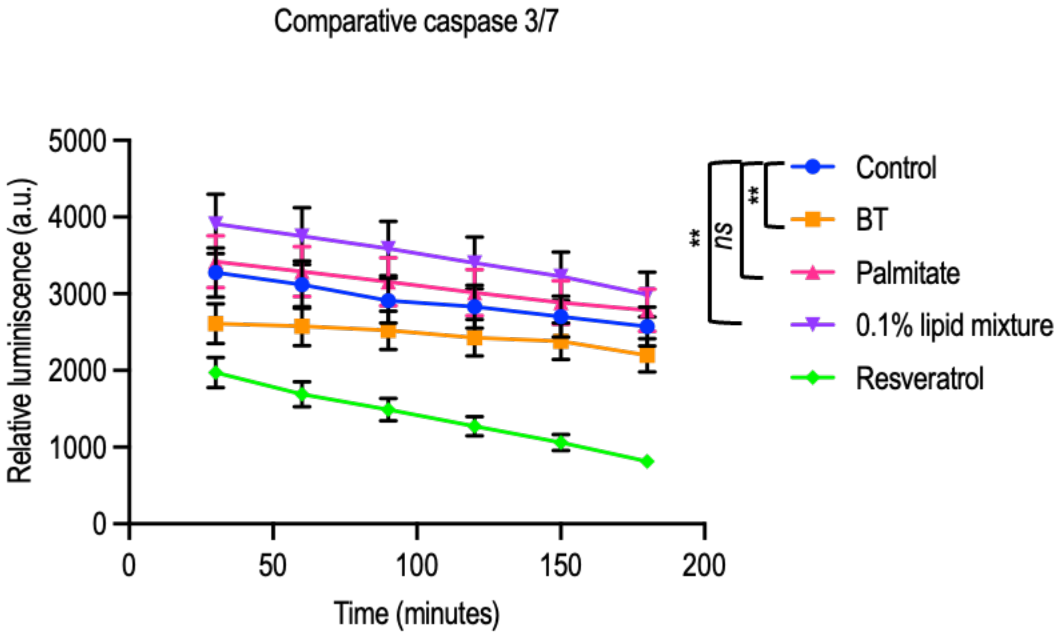


**Figure S6.** A comparative apoptosis profiling through the measurement of real time bioluminescence of caspase 3/7 activities in perfused cardiac cells treated with butyrate and different grade of fatty acid include- long chain fatty acid (palmitate), lipid mixture (0.1%) and standard control, resveratrol are depicted and quantified compared to control. All the BT treatment data were quantified and plotted based on three to five independent experiments and compared to control. The bar represents the mean ± SD (standard deviation). Statistical analyses were conducted using GraphPad Prism Version 10.3.1 (464), and significance level are denoted as $^{**}p<0.01$, based on Student's t-test. 'ns' denotes no significance.
